# Dynamical Regimes of Excitable Cells under Pulsed Electric Fields: From Excitation to Electroporation

**DOI:** 10.64898/2026.10.26.761774

**Authors:** Md Rashid Ali Faridi, Rohit Manchanda, Rochish M. Thaokar

## Abstract

Nanosecond pulse electric fields (nsPEFs) are an established treatment modality for cancer and have emerged as a potential approach for deep brain stimulation and cardiac defibrillation. The concerns in these two applications are at odds. While pulsed-field electroporation applications aim to minimise excitation of surrounding neurons, therapies based on neuronal stimulation demand minimal electroporation of neurons. Although nsPEFs offer various advantages, such as minimising power input and suppressing electrode reactions, they require strong electric fields to reach both the excitation and electroporation thresholds, so maintaining a safety gap between the two is crucial. Because these coupled processes unfold on time scales ranging from nanoseconds to milliseconds and cannot be resolved simultaneously in experiments, a computational approach is used. This study examines the interplay between excitation and electroporation within the Hodgkin-Huxley (HH) framework for a planar two-bilayer model coupled with the Asymptotic Smoluchowski Equation for electroporation, and solves for transmembrane potentials (TMPs), pore statistics and pore size evolution. The model can predict the role of electroporation in modulating neuronal responses and qualitatively explain key experimental observations in the literature. These include excitation with a single action potential (SAP), repetitive spiking (RS), and depolarisation block (DB), and are explained here with detailed mechanisms, including the subtle effects of strong electric fields and electroporating currents on ion-channel dynamics. Importantly, our work establishes a parameter space for strength-duration curves in regimes of safe excitation without electroporation, and vice versa. For a neuron with an equivalent RC time constant of 10 ns, pulses shorter than about 20 ns electroporate the membrane before they excite it, and the safety factor falls below 5% for 1-ns pulses. The work also expands into *µ*s and *m*s pulses, and delves into possible regimes that could lead to neuronal inviability due to irreversible electroporation or excessive fractional pore area.

**Author summary:** Very short, intense electrical pulses, lasting only billionths of a second, are used in cancer treatment, explored for brain stimulation, and proposed for cardiac defibrillation. Such pulses can make nerve cells fire electrical signals called action potentials, but they can also punch tiny holes (pores) in the cell membrane, a process called electroporation. Cancer treatment needs pores without stimulating nearby nerves, which causes pain and muscle contraction, whereas safe neurostimulation needs excitation without membrane damage. Although usually studied separately, these effects can occur together and influence one another. We developed a computational model that captures both, tracking how pores form, grow and alter the electrical behaviour of the neuron. We find that electroporation can reshape neuronal firing, producing repetitive spiking and depolarisation block as pulse strength increases, because pores open extra pathways for current that shift the balance of ionic currents. We map, for each pulse length, which stimulus strengths excite without porating, porate without exciting, or kill the cell. For very short pulses and small neurons, the safety margin between excitation and electroporation shrinks and can even reverse. Our findings provide a theoretical basis for designing more selective electrical stimulation protocols.

## 1 Introduction

The interaction between pulsed electric fields (PEFs) and excitable cells such as neurons and cardiomyocytes is of fundamental and translational importance. Two distinct yet increasingly overlapping domains—electroporation-based cancer treatment and electrical stimulation of excitable tissues—are being unified by the use of *ultrashort nanosecond pulsed electric fields* (nsPEFs). These modes of treatment offer new possibilities, but also raise fundamental questions about safety, efficacy, and the desired or undesired co-occurrence of membrane poration and cellular excitation.

In *oncology*, strong electric fields are applied to porate cell membranes for therapeutic purposes. Two major strategies include *irreversible electroporation* (IRE), which uses long and intense pulses to ablate malignant tissue (1), and *electrochemotherapy* (ECT), where short microsecond pulses reversibly porate membranes to facilitate drug uptake (2). A typical ECT protocol involves eight square pulses of 100 *µ*s and 0.6-1.5 kV/cm electric field range to induce reversible electroporation(ESOPE protocol) (3). Conventional IRE likewise employs microsecond pulses with electric fields of the order of kV/cm, but typically uses a substantially larger number of pulses and a greater overall electrical dose. More recently, nsPEFs have emerged as a promising alternative due to their ability to affect not only the plasma membrane but also internal organelles, including the nuclear membrane, mitochondria, and endoplasmic reticulum (4). These ultrashort pulses cause rapid capacitive charging of subcellular membranes and can induce poration. In such electroporation applications in oncology, excitable cells, including muscle and nerve cells, can be inadvertently affected, leading to pain, muscle contraction, and nerve damage (5).

In *neuroscience*, electrical pulses are used for therapeutic stimulation of excitable tissues. Applications such as deep brain stimulation (DBS), vagus nerve stimulation, and peripheral nerve modulation explore nsPEFs to reduce thermal effects, enhance depth penetration, and limit off-target activation (6). However, neuronal ion channels typically operate on millisecond timescales, posing questions about their ability to respond to nanosecond electric field pulses. The voltage required for nsPEF based stimulation reaches the limit of electroporating potentials, and an understanding of the effect of possible electroporation on the excitation and excitability of the excitable cells becomes critical.

A third area where nsPEFs are being explored is *cardiac defibrillation*. Defibrillation traditionally employs millisecond-scale pulses to depolarise heart tissue and terminate arrhythmias. Experimental results show that nsPEFs can excite isolated cardiomyocytes via membrane poration (7; 8). However, contradictory results have been reported at the tissue or organ level where nsPEFs show better efficacy than conventional pulses with reduced energy expenditure without any observable damage to the organ (9). Understanding whether defibrillation by nsPEFs is mediated primarily through electroporation or direct excitation is essential for future low-energy, non-thermal cardiac therapies.

Together, these trends raise several pressing questions with respect to the strength and duration of pulse:

- Generally *excitation* happens at a much lower stimulus without poration for millisecond conventional pulses. How do nsPEFs affect this safety gap between excitation and electroporation?
- Can *membrane poration* occur without triggering action potentials?
- What factors determine the threshold for poration vs. excitation across different pulse durations?
- How do ion channels and membrane conductance respond to ultrashort, high-intensity electric pulses?
- What is the size distribution of the pores formed during electroporation? How long do these pores live? This can have implications for ECT and gene delivery treatments.
- Can electroporation induce qualitative changes in the post-stimulus dynamics of excitable cells?

These questions are critical for developing safe and effective clinical protocols, especially when targeting mixed tissues of excitable (e.g., neurons, cardiomyocytes) and non-excitable (e.g., cancerous) cells. In cancer therapies, peripheral neurons may experience unintended excitation or damage during IRE or ECT. In neurostimulation, unwanted poration (10) may lead to cytotoxic effects.

The Pakhomov group (11) discusses three possible excitation mechanisms in the use of nsPEFs - excitation without electroporation, electroporation-induced excitation, and excitation independent of electroporation. Recent experiments on genetically engineered HEK cells reported diverse electrophysiological responses when subjected to pulsed fields (12). In general, there are contradictory results across various cell types, highlighting the lack of mechanistic understanding of such excitations under nsPEFs.

Much work has been done on modelling electroporation in biological cells, both analytical and computational, over the past four decades. The most fundamental approach in the continuum limit is to solve the Smoluchowski equation, in which the spatio-temporal pore density distribution is obtained as a solution of the appropriate partial differential equation (13; 14; 15; 16). The computational cost associated with solving the PDE is the limiting factor in this method (17). This changed with the landmark paper by Neu and Krassowska (17), which uses singular perturbation methods to transform the PDE into ODEs, in which pore density is evaluated as pore radii evolve. They are popularly known as the Asymptotic Smoluchowski Equations (ASE). A series of papers from 1999 to 2007 used various techniques to model electroporation in biological cells using the ASE (18; 19; 20; 21; 22; 23). The most noteworthy is the 2004 paper by Smith *et al*., where the cell is modelled as a membrane patch devoid of any spatial component (22). It computes the number of pores formed and their size evolution during and after the pulse, unlike earlier models (17; 18; 19; 20), that only compute pore number and do not account for pore-size dynamics. Krassowska *et al*. (23) added the geometrical complexity by solving it for a spherical cell using a finite-difference scheme and remains the most widely accepted continuum model of electroporation of a biological cell to date. Another line of computational studies for electroporation modelling often relies on finite element methods (FEMs) in COMSOL to evaluate field distributions, but rarely incorporates pore-size dynamics (24). The models discussed above have been used to study electroporation in non-excitable cells. The motivations stem from applications in ECT and gene transfection, where longer millisecond pulses are used.

The classical Hodgkin-Huxley model (HH-model) is a foundational model on which most of the computational modelling of excitable cells stands, and is still in use for modelling various excitable systems (25). In the context of PEFs, Boinagrov presented a HH-model-based strength-duration framework for single cells, challenging conventional excitation thresholds and emphasising the role of capacitive dynamics with extracellular stimulation (26). Subsequently few other authors (27; 28; 29) have presented HH-type models in an attempt to look at nsPEF-based excitation in neurons. These works do not address simultaneous electroporation modelling of neuronal cells under nsPEF stimulation.

There have been efforts to model electroporating neurons, notably by R.P. Joshi’s group, who modeled the incapacitation of neural traffic by conduction block of APs (30; 31). It solves an HH-type cable model coupled with Smoluchowski’s equation for cylindrical nerves, for both unmyelinated and myelinated nerves, for ultrashort pulses. It, however, does not delve into the excitation dynamics and associated mechanism. Another model incorporated an ASE electroporation model in the HH-model; however, their circuital model of the cell is oversimplified, and dynamic pore size evolution is missing (32; 33; 34). The work of Rems group on engineered HEK cells, also provided a computational model to explain the spiking responses in their experiments on genetically engineered HEK cells (12). This coarse-grained model uses FEM-based COMSOL Multiphysics solver to find the electric field in the domain, and excitability is captured by appropriate boundary conditions. An empirical relation is used to model electroporation, instead of solving ASE. Pore statistics and the size dynamics are therefore not provided in their analysis. While their model explains the widening of AP at higher stimulus strength, it fails to capture repetitive spikes and the spike latency as the stimulus strength increases.

In the present study, the ASE is incorporated into a space-clamped Hodgkin-Huxley-type neuron model to investigate the effects of excitation and poration across a wide range of pulse durations and intensities. The objectives set forth are:

1. Develop a space-clamped HH-type neuron model that incorporates electroporation by solving full ASE, and providing complete pore size statistics and dynamics.
2. Map the strength-duration response space and identify domains of:

- Excitation without poration
- Poration without excitation
- Excitation with poration
- Inactivity

3. Provide a mechanistic understanding of the responses observed in the model and possible corroboration with experiments

**Table 1.**
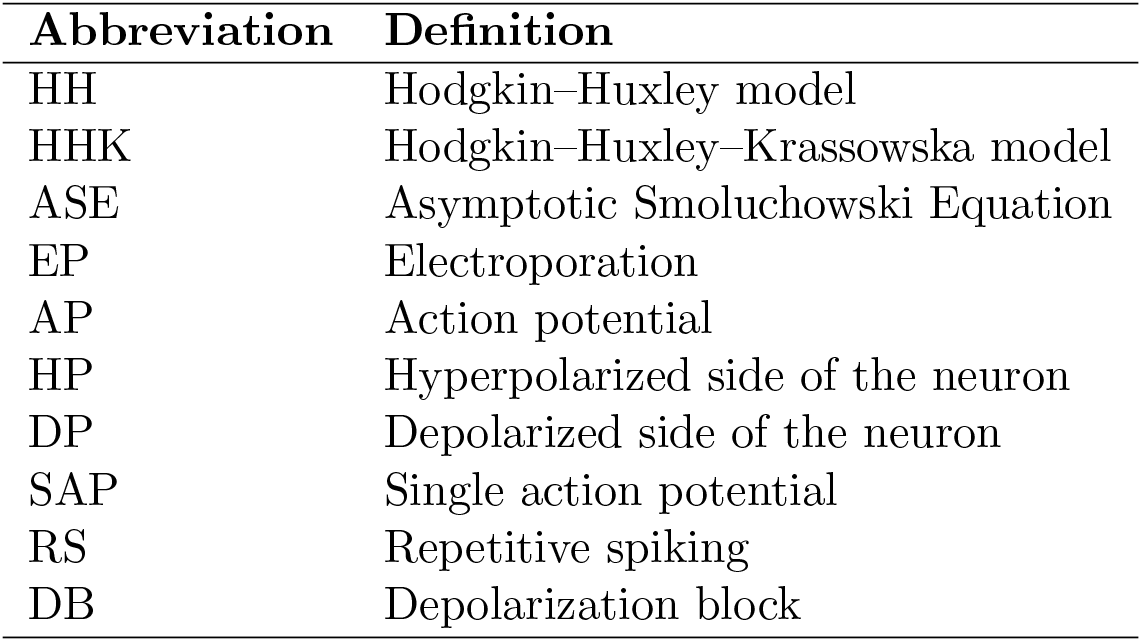
Abbreviations used throughout the manuscript.

This modelling effort also aims to clarify the mechanistic boundaries between reversible/irreversible electroporation and action potential generation. The findings may guide the design of tunable nsPEF waveforms for *excitation-free poration* or *poration-free excitation*, with translational relevance to cancer therapy, neurostimulation, and cardiac defibrillation.

## 2 Methodology

A typical neuron is considered, in which, instead of the exact neuronal geometry, the system is approximated using a planar cell model with two polarized sides of the neuron represented as two parallel bilayers, separated by an intracellular fluid, providing a minimal model (see Fig. 1). This approach simplifies geometry-related complexities, focusing on a bilayer system instead, to facilitate the understanding of excitation and electroporation processes. The membrane corresponding to the positive electrode (anode) is termed as hyperpolarized membrane (HP), while the other side, where the negative electrode (cathode) causes depolarisation, is termed as depolarized membrane (DP). The two sides are separated by cytoplasmic resistance, *R*. The Hodgkin-Huxley (HH) model is adapted with an additional shunt conductance, *g*_*ep*_, electrically parallel to the standard ion channel conductances in order to represent electroporation. The shunt conductance varies during and after stimulation, depending on the extent of membrane poration, including the number and size of electropores, as well as their subsequent closure and resealing.

**Fig 1.**
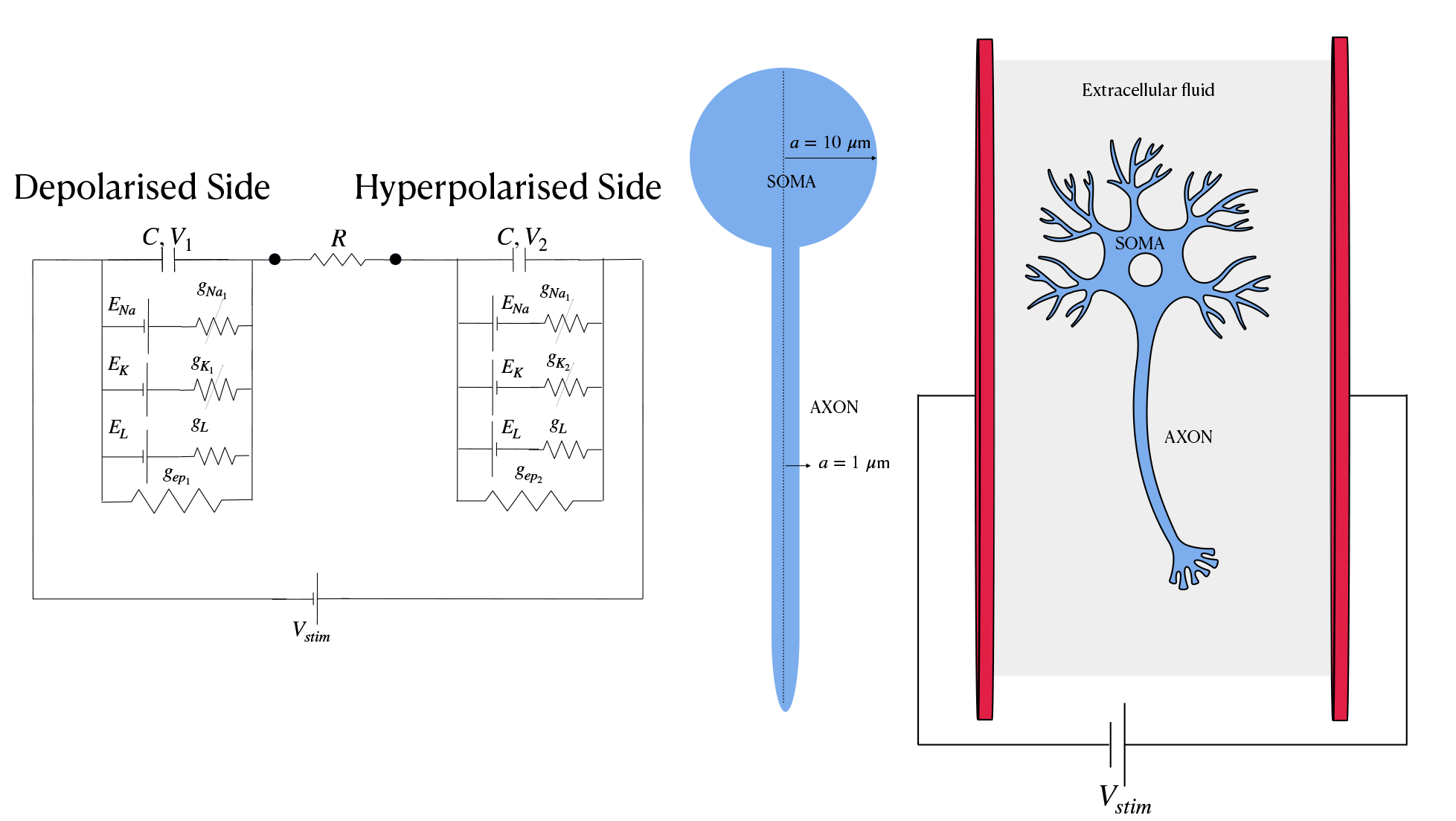
Schematic representation of stimulation of a planar two-bilayer model of a neuron, as a minimal model for a neuron, and its circuital representation.

### 2.1 Governing Equations

The transmembrane potentials, *V*_1_ and *V*_2_, (defined as the potential across the membrane, measured inside (intracellular) *minus* outside (extracellular)) correspond to the depolarized (DP) and hyperpolarized (HP) sides of the membrane, respectively. They are governed by the following coupled equations derived using the Kirchhoff’s law:

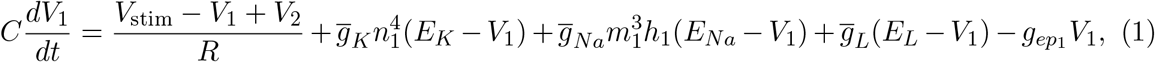

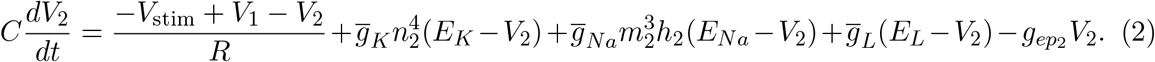

Here, *C* is the membrane capacitance, *V*_stim_ denotes the external voltage stimulus. 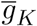] and 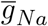] represent the maximal conductances for potassium and sodium ions, respectively. *E*_*K*_, *E*_*Na*_, and *E*_*L*_ are the Nernst (reversal) potentials for potassium, sodium, and leak currents (see table 2 for details of the various symbols used here and later). The electrical response of the cell, as represented by the circuit shown in Fig.1, is characterized by an equivalent *RC* time constant. It is related to the cell-charging time, *τ*_*p*_, through *RC* = 2*τ*_*p*_, where *τ*_*p*_ denotes the characteristic time for cell charging due to an externally applied electric field.

**Table 2.** Model parameters and initial conditions used in simulations for HH modeling.

| Symbol | Description | Value | Unit |
| --- | --- | --- | --- |
| $E_{Na}$ | Sodium Nernst potential | 45 | mV |
| $E_K$ | Potassium Nernst potential | -82 | mV |
| $E_L$ | Reversal potential associated with leak current | -59 | mV |
| $V_1(0)$ | Initial potential at DP side | -70 | mV |
| $V_2(0)$ | Initial potential at HP side | -70 | mV |
| $C$ | Membrane capacitance | 1.0 | $\mu\text{F}/\text{cm}^2$ |
| $R$ | Cytoplasmic resistance | 10 and 1000 | $\text{m}\Omega\cdot\text{cm}^2$ |
| RMP | Resting membrane potential | -70 | mV |
| $\bar{g}_{Na}$ | Maximal sodium conductance | 120 | $\text{mS}/\text{cm}^2$ |
| $\bar{g}_K$ | Maximal potassium conductance | 36 | $\text{mS}/\text{cm}^2$ |
| $\bar{g}_L$ | Leakage conductance | 0.3 | $\text{mS}/\text{cm}^2$ |
| $m_{10}, m_{20}$ | Initial activation variable of Na-channel | 0.053 | - |
| $n_{10}, n_{20}$ | Initial activation variable of K-channel | 0.317 | - |
| $h_{10}, h_{20}$ | Initial inactivation variable of Na-channel | 0.596 | - |

The ion channel dynamics follow standard Hodgkin-Huxley (HH) gating variables, which are updated based on the transmembrane potentials (TMP) *V*_1_ and *V*_2_.

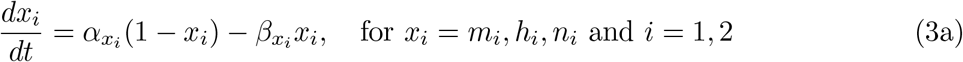

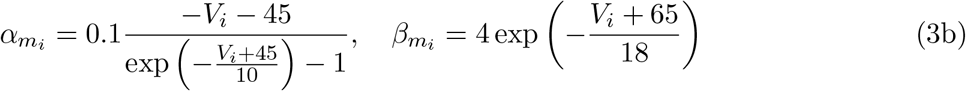

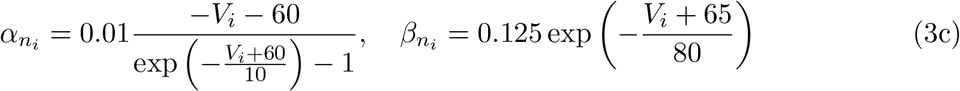

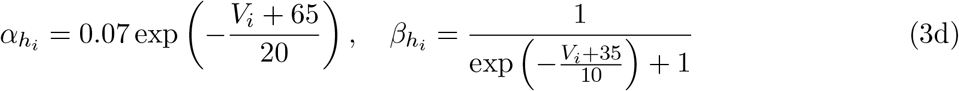

Here, *m* and *h* are the activation and inactivation gating variables for the voltage-dependent sodium channel, while *n* is the activation variable for the voltage-dependent potassium channel. *α* and *β* are the corresponding rate constants and depend upon the respective TMPs of the two bilayers. The subscript *i* = 1 refers to the DP side of the membrane, and *i* = 2 refers to the HP side.

While ionic currents depend on their respective Nernst potentials, the electroporation-induced (shunt) current does not, due to the larger radius of the electropores (approximately 0.8 nm and larger). They lose ion selectivity, allowing multiple ionic species to permeate simultaneously. Accordingly, the shunt current is modeled with a reversal potential of 0 mV. The electroporation, for each of the bilayers, itself is modeled using Krassowska’s Asymptotic Smoluchowski Equation (ASE), where the kinetics of the formation of these pores is described the equation,

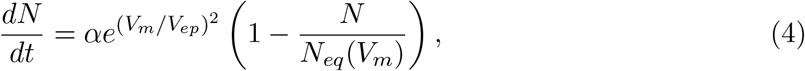

where *N* is the pore density, *V*_*m*_ is the TMP, *V*_*ep*_ is the characteristic voltage of electroporation ∼ 258 mV, and *α* is the creation rate coefficients. *N*_*eq*_ is the equilibrium pore density at a given TMP, *V*_*m*_.

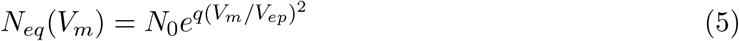

*N*_0_ is the equilibrium pore density at zero TMP and *q* = (*r*_*m*_*/r*_∗_)^2^. The definitions, expressions and values of parameters such as *w*_*o*_, *η*, etc. are provided in tables 2 and 3. The ASE and other equations presented here are detailed in the Appendix A. The shunt conductance *g*_*ep*_ depends on pore formation, growth, and resealing dynamics, modeled using

**Table 3.** Model parameters and initial conditions used in simulations for electroporation modeling. Parameters are taken from Smith *et al*. (22), except for *w*_0_ and *η*_0_, which are taken from Salimi *et al*. (37).

| Symbol | Description | Value | Unit |
| --- | --- | --- | --- |
| $\alpha$ | Creation rate coefficient | $1 \times 10^9$ | $\text{m}^2/\text{s}$ |
| $V_{ep}$ | Characteristic voltage of electroporation | 258 | mV |
| $N_{10}, N_{20}$ | Equilibrium pore density at TMP = 0 mV | $1.5 \times 10^9$ | pores/$\text{m}^2$ |
| $r_*$ | Minimum radius of conducting pores | 0.51 | nm |
| $r_m$ | Radius corresponding to energy minima at TMP = 0 mV | 0.8 | nm |
| $D$ | Diffusion coefficient for pore radii | $5 \times 10^{-14}$ | $\text{m}^2/\text{s}$ |
| $T$ | Absolute temperature | 310 | K |
| $\beta$ | Steric repulsion energy | $1.4 \times 10^{-19}$ | J |
| $\gamma$ | Edge energy | $1.8 \times 10^{-11}$ | J/m |
| $\sigma_0$ | Tension of the bilayer without pores | $1 \times 10^{-3}$ | J/$\text{m}^2$ |
| $\sigma'$ | Tension at water-hydrocarbon interface | $2 \times 10^{-2}$ | J/$\text{m}^2$ |
| $A_1, A_2$ | Membrane bilayer area at each end ($A/2$) | $6.3 \times 10^{-10}$ | $\text{m}^2$ |
| $F_{max}$ | Maximum electric force for TMP = 1 V | $0.7 \times 10^{-9}$ | N/$\text{V}^2$ |
| $r_h$ | Constant in Eqn.A6 (see Section S1.1 in the SI) | 0.97 | nm |
| $r_t$ | Constant in Eqn.A6 (see Section S1.1 in the SI) | 0.31 | nm |
| $d_m$ | Membrane thickness | 5 | nm |
| $\sigma_p$ | Conductivity of the extracellular fluid | 1.2 | S/m |
| $q_e$ | Electronic charge of an electron | $1.65 \times 10^{-19}$ | C |
| $k_b$ | Boltzman constant | $1.380649 \times 10^{-23}$ | J/K |
| $w_0$ | Pore energy barrier | 3.2 | - |
| $\eta_0$ | Relative entrance length of pores | 0.15 | - |

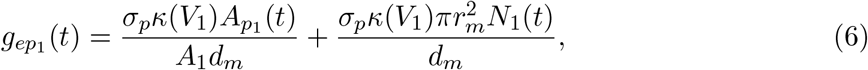

and,

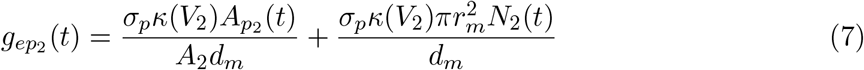

where,

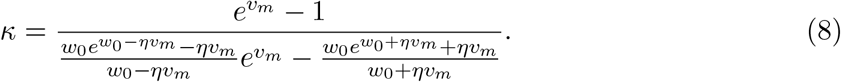

Here, *N* (*t*) denotes the pore density, and *A*_*p*_(*t*) is the total area occupied by the electropores, computed as 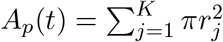*]. K* is the number of large pores, and *r*_*j*_(*t*) is the dynamic radius of the *j*^th^ pore. The function *κ*(*V* ) is a correction factor that captures the voltage-dependent energy barrier for ion flow through the nanopores (35) (also detailed in the Section S2 of the SI). These quantities evolve dynamically during and after the pulse, and are calculated separately for both sides of the membrane.

The constant *σ*_*p*_ denotes the conductivity of the liquid occupying the pore, and *d*_*m*_ represents the membrane thickness. *v*_*m*_ = *q*_*e*_*V*_*m*_*/k*_*b*_*T* is the non-dimensionalized transmembrane potential (TMP), and *q*_*e*_ is the electronic charge. For a spherical cell with radius *a* = 10 *µ*m, the total bilayer lipid area is approximately 1.26 × 10^−9^ m^2^. This area is equally divided between the depolarized (DP) and hyperpolarized (HP) sides of the membrane, as a simplifying assumption consistent with the current modeling framework.

#### 2.1.1 Relevant Biophysical Timescales

Solving Eqn.1 and Eqn.2 without taking into account the ion-channel dynamics, gives an estimate of *V*_1_ and *V*_2_ during the pulse duration. TMP at DP side estimates as 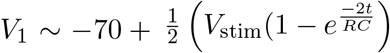], while TMP at HP side is 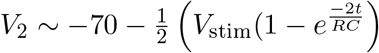].

Pore creation time can be estimated by assuming *N << N*_*eq*_ in Eqn.4. This gives:

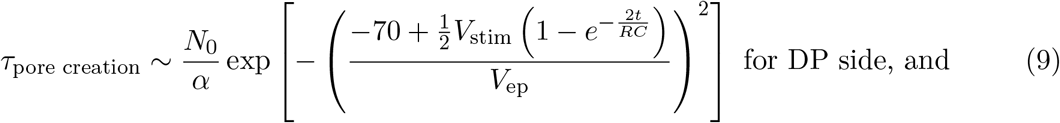

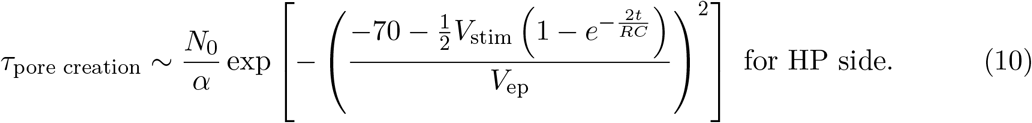

For a pulse of *τ* = 10 ns and *V*_stim_ = 2530 mV in a cell of *RC* = 10^−8^ s, pore creation time, for *τ* ∼ *τ*_*p*_ approximates to 200 ns and 2 ns at DP and HP sides respectively.

### 2.2 Numerical Strategy and Modeling Considerations

The model described by Eqns.1 - 8, and Eqns.A3 - A6 constitutes the Hodgkin-Huxley-Krassowska model (HHK), and is implemented using MATLAB R2023a (36), and executed on a MacBook Pro with an Apple M2 processor. The numerical details are provided in the Appendix B and are only briefly sketched here (see the schematic of the numerical implementation in Fig.S8). The classic ASE model, implemented here for both the HP and DP side bilayers (22), is modified by conducting simulations over *4* different sub-time spans. A monopolar rectangular pulse is given at *t* = 5 ms. All the ion-channel parameters are equilibrated over this time upto *t* = 5ms (*t*_1_). The ultrashort pulse of high intensity, typically, is on the order of ns to *µ*s (*τ* ). The pulsing can result in the formation and evolution of conducting (hydrophilic) pores. This happens over a time scale of a few *µ*s to ns (*τ*_pore creation_). However, excitation or firing of AP in neurons happens over few *milliseconds*. Numerically this is implemented as a sub-time span *t*_2_ which is the duration of the pulse, while the sub-span *t*_3_ equals *RC* of the cell, the timescale over which the TMP relaxes due to the resistivity of the cytoplasm. The magnitude of the membrane potential at both ends increases during the pulse, and the recovery post-pulse to the resting potential occurs over a few *time constants* of the cell. The combined period of *t*_2_ + *t*_3_ is the phase in which pores form and evolve, thereby encompassing time scales *τ* and *τ*_pore creation_. This phase is called *pore launching phase*. Resealing of these hydrophilic pores happens through their conversion into hydrophobic (non-conducting) pores and eventual annihilation by the lipid fluctuations. This occurs on a time scale of a few *seconds* (*t*_4_), termed as *pore absorbing phase*. Each of these subdivisions require a distinct solver configuration. The model solves for pore density *N* (*t*) through populations of small and large pores, and tracks the radii distribution (*r*_*j*_) on both the DP and HP side bilayers, as well as the pore closure, through the now well established and accepted numerical protocols listed elsewhere (22). The entire protocol is elaborated out in the Appendix B.

Neuronal dynamics specificity in the numerical algorithm for electroporation occurs during the *t*_2_-phase, when ultrashort pulses de-activate potassium channels and de-inactivate sodium channels at the HP side. The large negative TMP attained at the hyperpolarized side during nsPEF stimulation causes immediate deactivation of K-channels and de-inactivation of Na-channels on that side. The resulting decrease in potassium efflux, combined with an increase in sodium influx, continues to depolarize the membrane. As the system transitions into the *t*_3_-phase, the TMP begins to relax, and the gating variables *n*_2_ and *h*_2_ continue to evolve based on both the TMP and the injected current, during and after the stimulus. Capturing these rapid transients in the ion-channel variables, necessitates careful selection of the time step (Δ*t*) during integration in the pore launching phase.

## 3 Results

We present results for the equivalent *RC* time constant of 10^−8^ s (10 ns), that could correspond to a typical neuron with axon of radius of 1 − 2 *µ*m, and ionic conductivities of the intra-extra-ceullar fluids of the order ∼ 1 S/m (38), with specific membrane capacitance, corresponding to 1 *µ*F/cm^2^. We also present results, predominantly in the SI (Section S12 and Fig.S18), for *RC* = 10^−6^ s (1 *µ*s), corresponding to retinal ganglion, with radius of 5 *µ*m but conductances of the fluids of the order ∼ 0.0126 S/m (39). This range then covers neurons with a soma of radius of 10 − 20 *µ*m, and ionic conductivities of the intra-extra-cellular fluids of the order ∼ 1 S/m, which gives cell charging time (*τ*_*p*_) from 5 ns - 500 ns. It corresponds to the equivalent *RC*-range from 10 ns - 1 *µ*s (*RC* = 2*τ*_*p*_). Unless otherwise stated, all simulations and results are presented in terms of the equivalent *RC* time constant.

### 3.1 Nanosecond pulse electric field shows different excitation responses such as single action potential, repetitive spiking and depolarisation block

The model excitable cell, when subjected to a single nanosecond pulse electric field (nsPEF), shows a variety of responses that are sensitively dependent on the extent of electroporation and the associated change in the membrane conductance. A brief description of these responses is presented here using a representative 100 ns pulse (see Fig.2, note *RC* = 10^−8^ s unless otherwise specified)

1. A **Single action potential (SAP)** is observed with or without electroporation on the membranes, typically at lower *V*_*stim*_ values. Fig.2(a) shows a single action potential firing for *V*_stim_ = 1557 mV. Electroporation typically happens when TMP exceeds 1 V. Fig.2(a) shows that the TMP of neither side exceeds 1 V, as seen in the figure inset for the given stimulus. It indicates that the observed excitation is not due to electroporation and that the stimulus is below the electroporation threshold. Mechanisms of excitation (SAP) using representative pulse durations of 10 *µ*s, 1 *µ*s, and 1 ns at their respective excitation thresholds (HHK) are detailed in the Section S5 in the SI for neurons of *RC* = 10^−8^ s.
2. **Repetitive spiking (RS)** is a behaviour shown by neurons where they fire in quick succession, often due to some external stimulus. As *V*_stim_ is increased, the model neuron shows repetitive spiking due to mild electroporation, and is typically observed for relatively higher stimuli than the excitation threshold. This repetitive spiking is transient in nature. It stops after some time, and the cell reaches the resting membrane potential (RMP) of -70 mV. Fig.2(b) shows a repetitive spiking behavior for *V*_stim_ = 2300 mV.
3. **Depolarisation block (DB)** occurs when a cell fires an action potential but fails to reach its initial resting membrane potential, instead plateauing for a long time at a significantly de-polarized state. This state renders the cell unexcitable. To demonstrate this feature, *V*_stim_ of 2479 mV is applied (see Fig.2(c)). The depolarisation block features extensive electroporation at both ends of the model neuron. At the given field, the effect is transient and the TMP eventually goes to RMP.

These responses indicate that electroporation, which is the formation of non-selective non-specific pores on the cell membrane, in the context of excitable cells, regulates excitation responses - the SAP, the repetitive spiking and the depolarisation block. The subsequent section provides a detailed discussion of this interplay.

**Fig 2.**
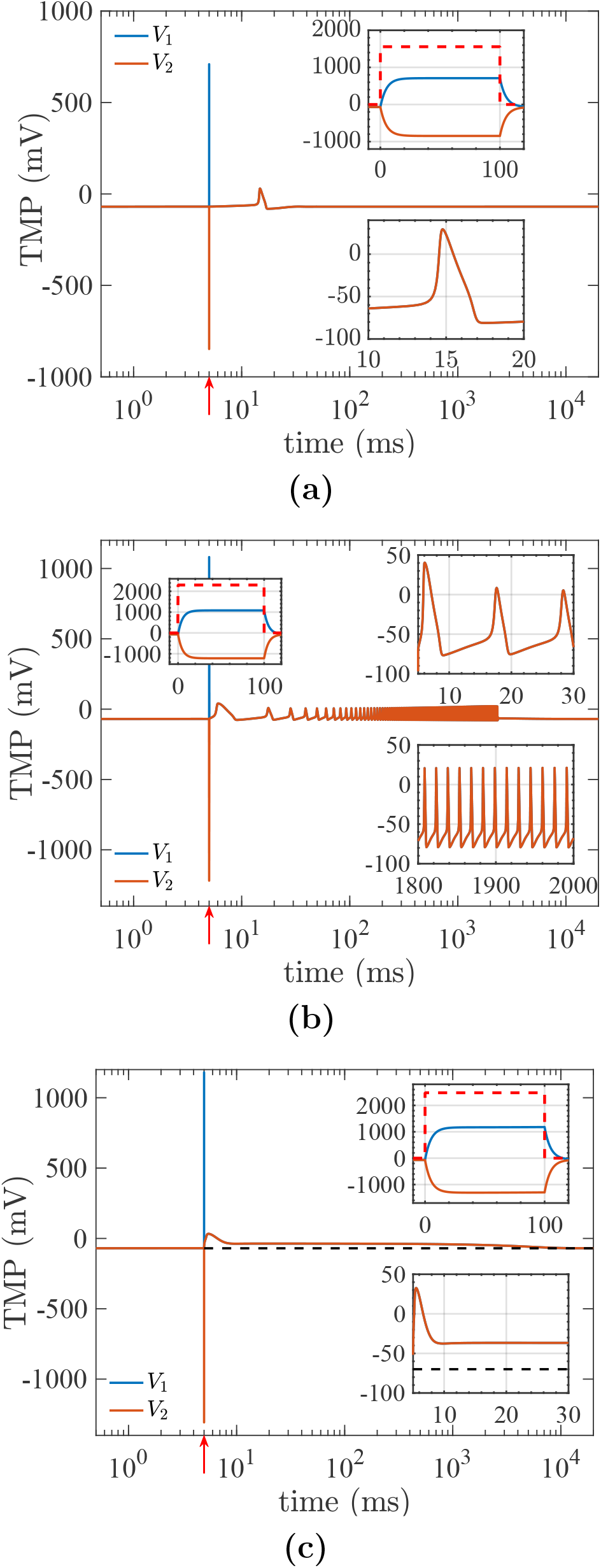
TMP dynamics in response to a 100 ns pulse applied at *t* = 5 ms. (a) Single action potential (SAP) at *V*_stim_ = 1557 mV; (b) repetitive spiking (RS) at *V*_stim_ = 2300 mV; (c) depolarization block (DB) at *V*_stim_ = 2479 mV. Insets show the stimulus waveform of 100 ns rectangular monopolar pulse (**– –**) with time in ns, and the corresponding charging– discharging dynamics of the depolarized and hyperpolarized membrane regions on the ns-time-axis. Another inset also shows millisecond-scale action potentials on the ms time-axis. In Fig.2(b), an additional inset highlights sustained uniform firing in RS over 1800–2000 ms. The stimulus onset at *t* = 5 ms is indicated by the upward red arrow (↑). For the inset showing the rectangular stimulus pulse, time is reset to t=0 at the stimulus onset to better visualise the TMP dynamics during pulse application.

### 3.2 nsPEF vs. longer pulses: Strength-duration picture for *RC* = 10^−8^ s neuron

A detailed picture of all possible excitation responses in the form of strength-duration (SD) curves, highlighting regimes of different responses in a regime map, is presented in Fig.3. Thresholds for different responses are defined as follows:

- **Excitation threshold**: Minimum *V*_stim_ that elicits a single action potential in the cell. One calculated using only the Hodgkin-Huxley model is abbreviated as HH, while the threshold computed using the Hodgkin-Huxley model incorporating the ASE model of electroporation by Krassowska *et al*. is abbreviated as HHK (22).
- **Electroporation threshold**: *V*_stim_ that shows a single pore formation at any of the two sides of the cell.
- **Repetitive spiking threshold**: When the application of a pulse fires two or more action potentials, it is termed repetitive spiking, and the minimum *V*_stim_ that can trigger such a behaviour is considered its threshold.
- **Depolarisation block threshold**: Minimum *V*_stim_ for which the cell exhibits depolarisation block. As a transient feature develops due to electroporation, the response qualifies as a depolarisation block if TMP stays above -40 mV and unable to reset back to its RMP.
- **Cell death threshold**: The voltages showing cell destruction either due to irreversible electroporation (IRE) or supra-electroporation.

Two novel features that come out of the SD curve in Fig.3 are:

1. The richness of the excitation responses, such as - SAP, repetitive spiking and depolarisation block, lies to the left, i.e., in the nsPEF range, and it ceases for longer pulses beyond 2*µ*s, whereafter only SAPs are seen in the *µ*sPEF and msPEF ranges (see Fig.3(a) for the complete SD curve for pulse durations ranging from 1-ns to 1-ms. Fig.3(b) shows the SD curve in the nsPEF regime for pulse durations 10-ns to 1-*µ*s.)
2. Neurons with *RC* = 10^−8^ s and pulse width of 10 ns and below, show electroporation even below the excitation threshold as shown in Fig.3(c).

The excitation behavior could be sensitively affected by electroporation; hence, electroporation thresholds are also presented. Together, these delineate the excitation regimes in the SD curve: electroporation without excitation, SAP with electroporation, repetitive spiking, and depolarization block. For pulses in *µ*sPEF-msPEF range, there seems to be a healthy gap between low excitation thresholds and the high electroporation threshold; this gets dangerously narrower for shorter pulses wherein ultimately for pulses below 10 ns for *RC* = 10^−8^ s, the electroporation threshold becomes less than the excitation threshold, thereby indicating electroporation before the firing an AP. The possibility of smaller excitable cells getting electroporated before excitation using ultrashort pulses is thus demonstrated here.

The dependence of various response thresholds on the cell-charging time (*τ*_*p*_ = *RC/*2) is explored in the subsequent section. SD curves for a larger cell (*RC* = 10^−6^ s) are presented in Fig.S18 of the SI.

### 3.3 Single action potential, repetitive spiking and depolarisation block triggered by nsPEF: Interplay of excitation and electroporation

The excitation response of a single action potential could be without triggering electroporation (see the detailed discussion later in Section 3.4 and (26)). On the contrary, repetitive spiking and depolarisation block are observed in neurons using nsPEF, and result exclusively from electroporation at both ends of the neuron. The rapid creation of transient non-selective ion channels, known as pores, on the cell membrane raises the membrane conductivity. The membrane specific conductance, on account of electroporation, is denoted by *g*_*ep*_, and is calculated for both the sides (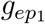] - DP side, 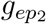] - HP side). Computations for a representative 100-ns pulse in a neuron of *RC* = 10^−8^ s are presented here in Fig.4 and in Table 4 to demonstrate the change in the membrane conductivity incurred by the nsPEF. The membrane conductance due to electroporation (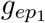] and 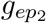] ) is illustrated along with pore statistics at thresholds for different responses. The subsequent analysis holds for other pulses in the nsPEF regime. A distinct feature of the plots *is the asymmetry in how the neuron experiences poration, i*.*e., the electroporation at the DP side is negligibly small compared to the HP side*. This is in accordance with the experimental findings that the hyperpolarised part of the cell experiences more poration than the depolarised side (40; 41), mainly on account of the negative resting potential. Some key observations from the Fig.4 are as follows. :

**Fig 3.**
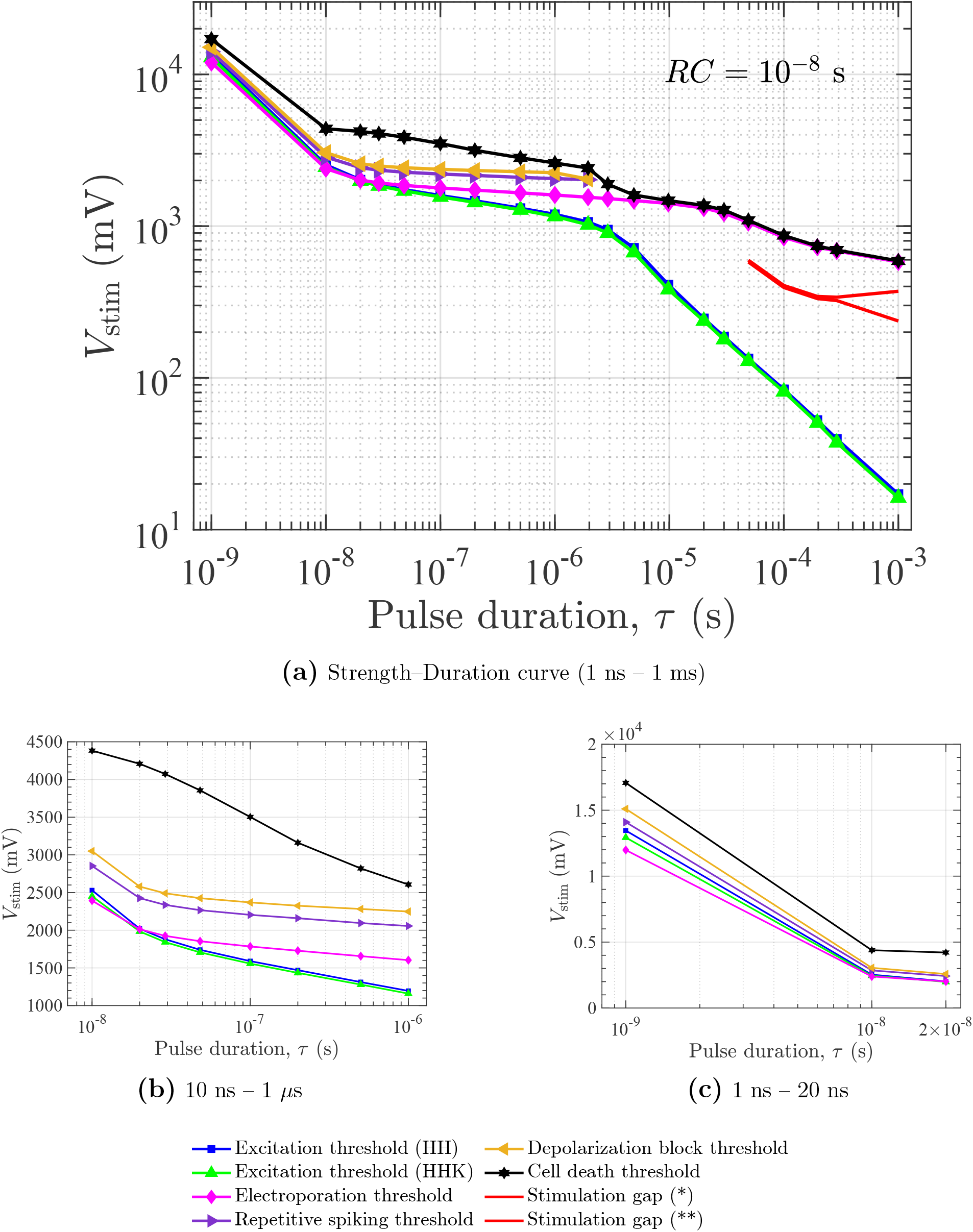
SD curves (*V*_stim_ vs pulse duration (*τ* )) for a neuron with *RC* = 10^−8^ s in the regimes a) 1 ns to 1 ms, b) nsPEF regime 10 ns – 1 *µ*s, c) 1 ns – 20 ns, which specifically highlights electroporation without excitation. Thresholds of excitation responses - single action potential, repetitive spiking and depolarisation block are provided alongside the electroporation threshold and cell death threshold. *Stimulation gap* is a small region in the ms-regime where there is no excitation, albeit the stimulus is above the excitation threshold (also see (26)). (^*^ indicates *lower threshold*, ^**^ indicates *upper threshold* in *Stimulation gap*).

1. Electroporation conductance increases with stimulus strength. The HP side conductance due to electroporation 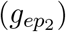 at applied stimulus of 2100 mV, 2300 mV, and 2479 mV reaches 0.6, 17, and 511 mS/cm^2^, respectively, by the end of the pulse. In contrast, the DP side shows a marginal increase in pore conductance due to electroporation, i.e., 0.08, 0.49, and 3.3 mS/cm^2^, respectively (compare figures 4(a),4(b),4(c)). It also supports the hypothesis that the nature of the neuronal excitation responses, i.e., single action potential, repetitive spiking and depolarisation block, depend primarily on the extent of electroporation.
2. A modest rise of 18% (*V*_stim_ changed from 2100 mV to 2479 mV) in stimulus strength shoots up the membrane conductance by 852-fold at the HP side, and 40-fold at the DP side, covering the spectrum of responses from SAP to repetitive spiking to depolarisation block (see Table 4).
3. The HP side is clearly more affected due to electroporation. It should be noted that on application of voltage, the HP side clearly builds a higher potential (in magnitude) than the DP side, since the membranes on either side have a resting potential of around −70 mV. The HP side’s sensitivity is explained by an order or two magnitude of pores more than the DP side (Fig. S14 in the SI). The results further support the assertion of asymmetric poration in neurons with both *RC* values, as the pore number statistics for *RC* = 10^−6^ s are also provided in the SI (Section S7).
4. The insets of Fig.4(a),4(b),4(c) highlight that the HP side conductance reaches its maximum value towards the end of the pulse. The membrane gets fully charged for nsPEFs that are larger than the membrane charging time (5-ns cell-charging time (*τ*_*p*_ = *RC/*2) vs pulse time *τ* = 100-ns). As mentioned in the Section 2.2, in the simulations, the pores are allowed to launch at regular intervals during the pulse. The sharp rise of membrane conductance towards the end of the pulse is the culmination of increment in number of electropores and their size growth. Pores of radii 1.2-1.5 nm are seen with a 100-ns pulse in *RC* = 10^−8^ s neuron (see *r*_*max*_ in Table 4). Details of the pore radius growth dynamics for the DP and HP sides are provided in the Section S7 of the SI, see Fig.S15.
5. Fig.3(a) shows the disappearance of repetitive spiking and depolarisation block in the *µ*sPEF and msPEF regime. Post *τ* = 1 *µs*, the regimes are restricted to well-separated electroporation and excitation thresholds. The electroporation does not lead to any further modification in excitation since it is immediately followed by cell death, making its discussion inconsequential. For example, a 100 *µ*s pulse at the maximum allowable stimulus of 867 mV could induce only 0.053 mS/cm^2^ membrane conductance. The depolarising current due to the nanopore would not drive the cell into repetitive spiking and depolarisation block. It explains the disappearance of repetitive spikes and depolarisation block for longer pulses. See Fig.4(d) for reference.

**Fig 4.**
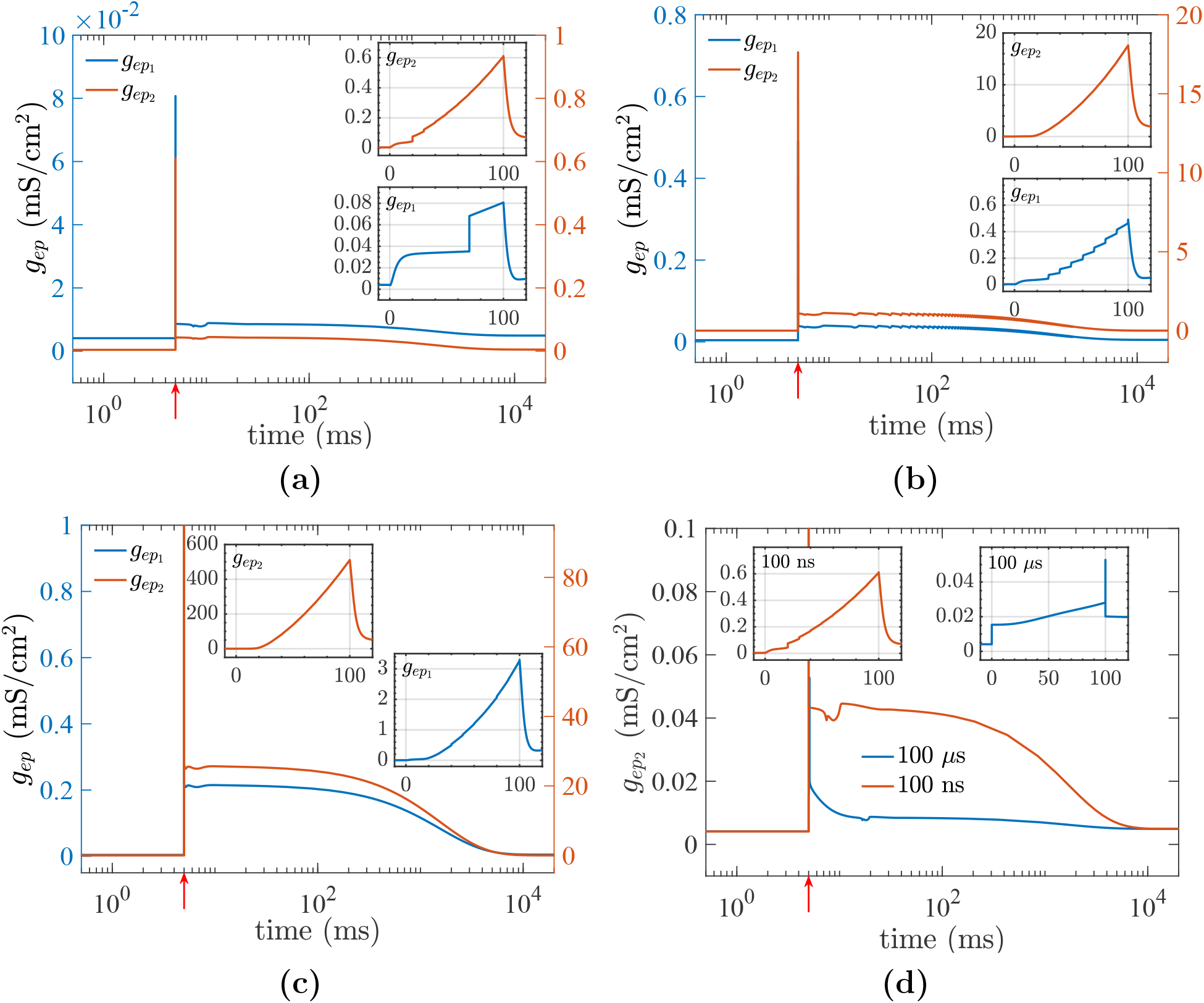
Pore conductance evolution for a 100 ns applied at *t* = 5 ms showing a) single action potential at *V*_stim_ = 2100 mV, b) repetitive spiking (RS) at *V*_stim_ = 2300 mV, c) depolarisation block (DB) at *V*_stim_ = 2479 mV (Left Y-axis - DP side, right Y-axis - HP side). d) HP side pore conductance evolution for 100 *µ*s and 100 ns pulse for *V*_stim_ = 867 mV and 2100 mV respectively corresponding to SAP with poration. The stimulus onset at *t* = 5 ms is indicated by the upward red arrow(↑). The inset illustrates the maximal rise in pore conductance during the pulse (time reset to *t* = 0 at pulse onset).

**Table 4.**
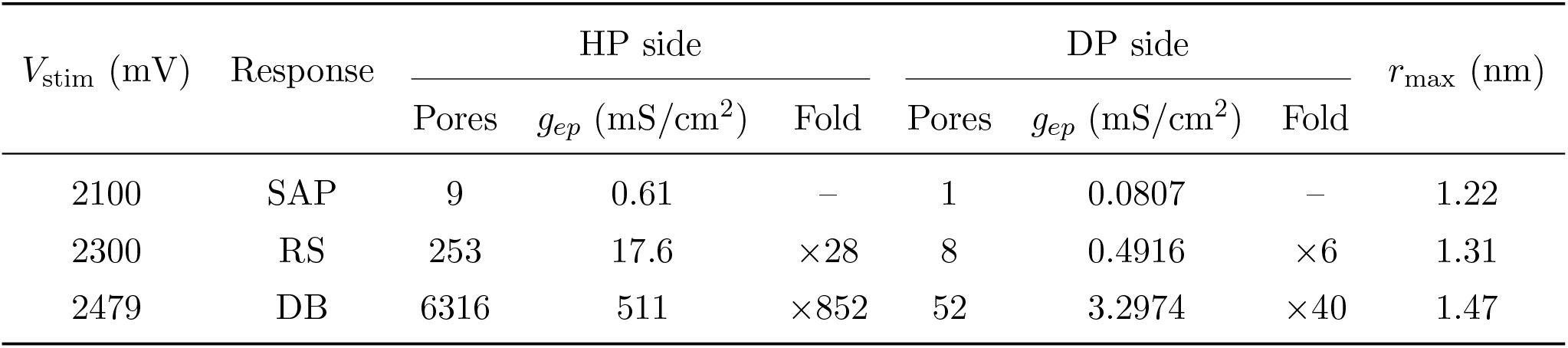
Electroporation-induced pore formation and conductance responses on the HP and DP sides for different stimulus strengths during a 100 ns pulse in a neuron with *RC* = 10^−8^ s. Fold changes are reported relative to the single action potential (SAP) stimulus level. SAP, RS, and DB denote single action potential, repetitive spiking, and depolarization block, respectively. *r*_max_ represents the maximum pore radius corresponding to a given *V*_stim_.

| $V_{\text{stim}}$ (mV) | Response | HP side | | | DP side | | | $r_{\text{max}}$ (nm) |
| --- | --- | --- | --- | --- | --- | --- | --- | --- |
| | | Pores | $g_{ep}$ (mS/cm<sup>2</sup>) | Fold | Pores | $g_{ep}$ (mS/cm<sup>2</sup>) | Fold | |
| 2100 | SAP | 9 | 0.61 | – | 1 | 0.0807 | – | 1.22 |
| 2300 | RS | 253 | 17.6 | $\times 28$ | 8 | 0.4916 | $\times 6$ | 1.31 |
| 2479 | DB | 6316 | 511 | $\times 852$ | 52 | 3.2974 | $\times 40$ | 1.47 |

### 3.4 Effects of electroporation on excitation dynamics

#### 3.4.1 Mechanism of neuronal excitation under nsPEF: Role of voltage-gated ion channels and their dynamics

To elucidate the effect of nanosecond pulses on the dynamics of voltage-gated ion channels and the corresponding effect of electroporation when the conditions induce electroporation, it is pertinent to discuss the excitation mechanism in an unporated neuron. A representative 100-ns pulse is considered as an example to illustrate the excitation mechanism in the model neuron below the electroporation threshold. The following features are noteworthy:

1. During the pulse, at the DP side, both the sodium channels and potassium channels respond minimally and see a marginal increase in conductance(see Fig.5(a), 5(c)) . It translates into a steady outward potassium current, with a minimal outward sodium current from the DP side 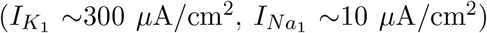. It should be noted that depolarising inward current is negative, while hyperpolarising outward current is positive, as per usual convention. Thus at the DP side, the state of the sodium and potassium ion channels does not change much.
2. At the HP side, the TMP quickly reaches -850 mV during the pulse (see inset Fig.2(a)). Note for a 100-ns pulse, the excitation threshold is 1557 mV, and for *τ >> τ*_*p*_, the TMP rises very sharply during the duration of the pulse. The sodium conductance 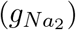 decreases much more rapidly (inset Fig.5(b)), than the potassium conductance 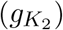 (inset Fig.5(d)) during the pulse. This is because 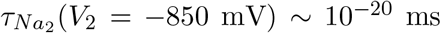], suggesting instantaneous dynamics, while *τ*_*K*_ (*V*_2_ = −850 mV) ∼ 10^−4^ ms (also see Table A1). The extremely small timescales, particularly ∼ 10^−20^ ms and below, should not be interpreted as characteristic times of a physical process; rather, they arise from the mathematical behavior of the Hodgkin–Huxley (HH) rate equations (Eqn.3) at large negative membrane potentials and indicate effectively instantaneous dynamics. Sodium channels on the HP side completely shut off, resulting in negligible inward depolarising current during the pulse (inset Fig.5(b), also exhibited in 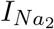] in the inset of Fig.5(e), also see the dynamics of the gating variables (*m, h* and *n*) on the DP and HP side in Fig.S12 and Fig.S13 in the SI). However, the potassium conductance drops from 0.4 to 0.2 mS/cm^2^ much slowly, causing a decaying inward depolarising potassium current during the pulse (inset Fig.5(d), also follow 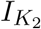] in the inset of Fig.5(f)).
3. Post-pulse relaxation of TMP brings down the DP side outward K-current, as seen in the upper inset of Fig.5(f) corresponding to the pulse duration time window (see 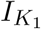] ). Meanwhile, recovery of the K-channel on the HP side occurs in milliseconds 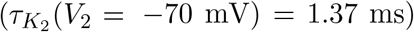. During this phase, the sodium current keeps creeping up from the HP side, causing a cumulative rise in the inward depolarising sodium current as seen in the lower inset of Fig.5(e) corresponding to the 4-18 ms time window (compare 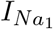] and 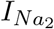] ). Clearly, the depolarising inward Na-current is slightly larger on the HP side. Note that this is facilitated by *h*_2_ ≈ 1 for the inactivation of Na-channels at the HP side. Hence, an AP is observed, primarily facilitated on the HP side by the fast deactivation gate of their K-channels during the pulse and their slow recovery after the pulse, thereby limiting the hyperpolarisation, and the simultaneous fast re-activation (opening) of the sodium channels on the HP side that were closed during the pulse, that induces inward depolarising current, resulting in an AP. This is referred to as K-dominated APs by Boinagrov *et al*.(26)(details are presented in the Sections S5 and S16.7, see Fig.S9). These differences in gating dynamics (*m, n*, and *h*) also manifest as the aforementioned K-dominated APs, primarily due to the asymmetry of time-constant functions for positive and negative TMPs, as seen on the DP and HP sides, respectively, during the pulse duration.
4. The higher the stimulus, the greater the deactivation of outward K-current at the HP side as seen in Fig.5(d) (also see inset therein), and thereby weaker the hyperpolarising current associated with the K-channels. Therefore, the net current is more depolarising (aided by opening of Na-channels at HP side) and early attainment of the minimum current required to fire the AP. This holds across different excitation regimes, including single action potentials (with and without electroporation), repetitive spiking, and depolarisation block (corresponding plots of ionic currents are shown in the Fig.S16 of the SI). Therefore, there is a reduction in spike latency with the increased stimulus (discussed later in the subsequent Sections 3.4.2 and 3.4.3).

A key question here is how incorporating the EP model affects the SD curve predicted by the HH model alone (26). The excitation thresholds predicted by the HHK model in the present study are always lower than the ones predicted by the HH model, *even for an unporated neuron*, and show a reduction up to 6% (Fig.S19 in the SI). The classical HH model does not solve for pore density *N* . Incorporating Krassowska’s electroporation model into the HH model yields the HHK model, with the addition of *g*_*ep*_*V* terms to account for pores forming stochastically due to lipid fluctuations at the cell membrane. Hence, *N* evolves, even when there is no electroporation and equilibrates to *N*_*eq*_ such that 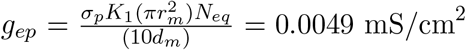]. This is negligible as compared to say *g*_*L*_ = 0.3 mS/cm^2^, but non-zero in non-electroporating regimes. This marginally increases the membrane conductance. Here, we term a non-electroporating system as one in which the number density of pores never exceeds *N*_*eq*_, and the pore size remains the equilibrium pore size, *r*_*eq*_=0.8 nm. This manifests as a slight rise in the conductance at the end of the equilibrating phase; otherwise, the ion-channels show the same dynamics during the pulse with a slight shift in their *m, n, h* and, *τ* values. Hence, the threshold in the HHK model is lower than that in the HH model.

#### 3.4.2 Electroporation alters features of action potential: Main features

The SD curve in Fig.3 suggests that for pulses below 10 ns for *RC* = 10^−8^ s, a neuron admits electroporation below the excitation threshold. We consider a 1-ns pulse to explore the effects of electroporation on the features associated with AP. Fig.6(a) compares the various features of the model computed using the HH (unporated) and HHK (porated) models for a pulse width *τ* = 1 ns, at various *V*_stim_, to clearly elucidate the effect of electroporation. Since the pulse duration *τ* = 1 ns, is smaller than membrane charging time (10 ns), hence the membranes are not fully charged during the pulse duration (see insets). The difference in the features of Aps for the porated and unporated neurons is distinct. To keep the comparison tractable, three key metrics are studied to quantify the differences in AP features computed using the unporated HH and HHK models: spike latency, the 50% recovery time from the peak (*T*50%), and the peak transmembrane potential (TMP) during the AP. As discussed earlier, the unportaed neuron only shows SAP for single pulses of increasing strength; however, the excitation response for the HHK model shows SAP, repetitive spike and depolarisation block. We first discuss the features of only the first AP after the pulse for all the regimes.

**Fig 5.**
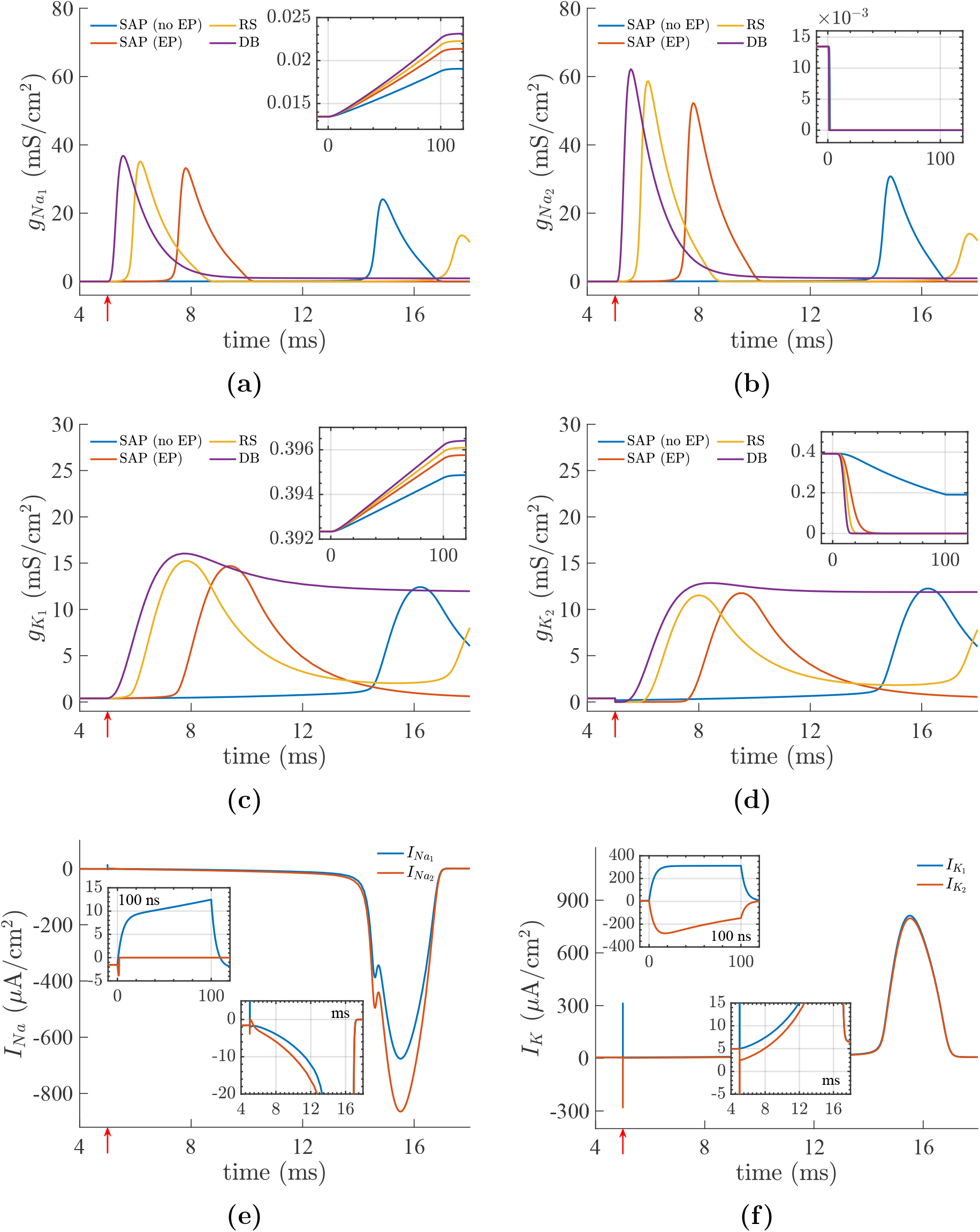
Na- and K-ion-channel conductance dynamics at the DP [(a), (c)] and HP [(b), (d)] sides are shown for four representative stimulation regimes: (i) single action potential without electroporation, SAP (no EP), at *V*_stim_ = 1557 mV (below EP threshold); (ii) single action potential with pore formation at both sides, SAP (EP), at *V*_stim_ = 2100 mV (above EP threshold); (iii) repetitive spiking (RS) at *V*_stim_ = 2300 mV; and (iv) depolarization block (DB) at *V*_stim_ = 2479 mV for a 100-ns pulse in *RC* = 10^−8^ s neuron. The insets enlarge the conductance dynamics during the pulse with the time axis reset to *t* = 0 at pulse onset for clarity. Panels (e) and (f) show the corresponding Na and K currents, respectively, for SAP (no EP) at *V*_stim_ = 1557 mV. Their upper insets highlight the current dynamics during the pulse (time reset to *t* = 0 at pulse onset), while the lower insets show the post-pulse current dynamics on the millisecond timescale leading to action potential generation, thereby delineating the differences in the Na and K ionic currents between the DP and HP sides. The stimulus onset at *t* = 5 ms is indicated by the upward red arrow(↑).

1. Fig.6(a) compares the action potential at a pulse strength of 13457 mV, the excitation threshold predicted using the unporated (HH) model. With the HHK model, the neuron fires AP and simultaneously gets porated, having six pores on the HP side. Spike latency is defined as the duration between the pulse onset and the peak of the first AP. The latency for the porated neuron (around 4ms), is clearly smaller than for the unporated neuron (10 ms) at the same *V*_stim_ = 13457 mV. This is often used as an indicator of poration.
2. The spike latency computed using HH and HHK shows a similar trend - monotonically decreasing with the increasing strength, in accordance with the experiments on excitable cells (12). For SAPs, this latency is around ten milliseconds to a few milliseconds. However, for repetitive spiking and depolarisation block, it is much lower. The spike latency saturates at 0.5 ms and 3.6 ms for APs computed using HHK and the unporated neuronal HH model (Fig.6(b)). A detailed discussion is presented in the subsequent section.
3. The time taken for the peak value of TMP (which the neuron attains post-early depolarisation phase during an action potential) to repolarise to 50% of its peak value is used to estimate action potential duration (APD) and is indicated as *T*_50%_. Fig.6(c) shows the effect of electroporation on *T*_50%_ in detail. The HHK model shows a larger *T*_50%_ than the HH model, indicating slower decay. It also validates the hypothesis of (12)- increased APD indicates electroporation of the cell membrane of neurons. As the extent of electroporation increases with *V*_stim_ and the response shows transitions of SAP to repetitive spikes and depolarisation block, so does the *T*_50%_ increase. Different gradients of the APD vs *V*_stim_ can be indicative of different excitation response types.
4. The peak transmembrane potential (TMP) during the AP, denoted as 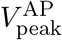], for the unporated HH model increases monotonically from +30 mV and then saturates around +39 mV (peak) for an unporated neuron. For the HHK model for porated neuron, the AP amplitude increases minimally from SAP to repetitive spike, and then drops sharply as the depolarisation block intensifies due to electroporation.(Fig.6(d)).

#### 3.4.3 Mechanistic insights into the role of electroporation in altering the features of action potential

Fig.4 and Table 4 establish that more pores are selectively formed at the HP side. The HP side sees immediate deactivation of K-channels followed by a subsequent slow recovery on cessation of the pulse. The slow buildup of the inward, depolarising Na-current from the HP side over this long recovery period allows the AP to fire. Pore formation, especially at the HP side, helps this cause by injecting a depolarising current. Near the excitation threshold for a 1-ns pulse with a 13457 mV stimulus, the cell already sees six small pores at the HP side. During the duration of the pulse and immediately after the pulse, the electroporation current through the pores is marginal (∼300 *µ*A/cm^2^ at HP side) compared to the stimulating current ∼ 10^9^ *µ*A/cm^2^ (stimulating current is mostly manifested as the displacement current of the membrane, details are provided in the SI (Section S9)). Electroporation intensifies as the pulse strength increases, thereby injecting more electroporation current inside the cytoplasm from the HP side. As the pores shrink, the membrane resealing happens over a few 100 s of milliseconds to seconds, pushing the decreasing current into the cell before it completely disappears from the cell membrane. Clearly, these features cannot be admitted by the simple HH model for an unporated neuron.

The sharp increase in the depolarising current due to the creation of pores during the pulse is stimulus-dependent. The greater the stimulus, the larger the number of pores created. However, they disappear slowly from the membrane. Hence, the observed difference between 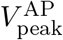 calculated using the unporated (HH) and porated (HHK) models. The electroporation current from the HP side is 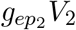]. Post stimulus, when *V*_2_ remains around the RMP, the pores disappear over a few seconds, allowing the depolarising current to gradually reduce, as the number of pores reduce, driving the system in this duration, through repetitive spiking as illustrated in Fig.6(e). The figure shows that the poration current remains predominantly depolarising (negative) at the HP side, corresponding to the depolarized TMP at the HP end, although the action potential oscillates between a hyperpolarised and depolarized state.

**Fig 6.**
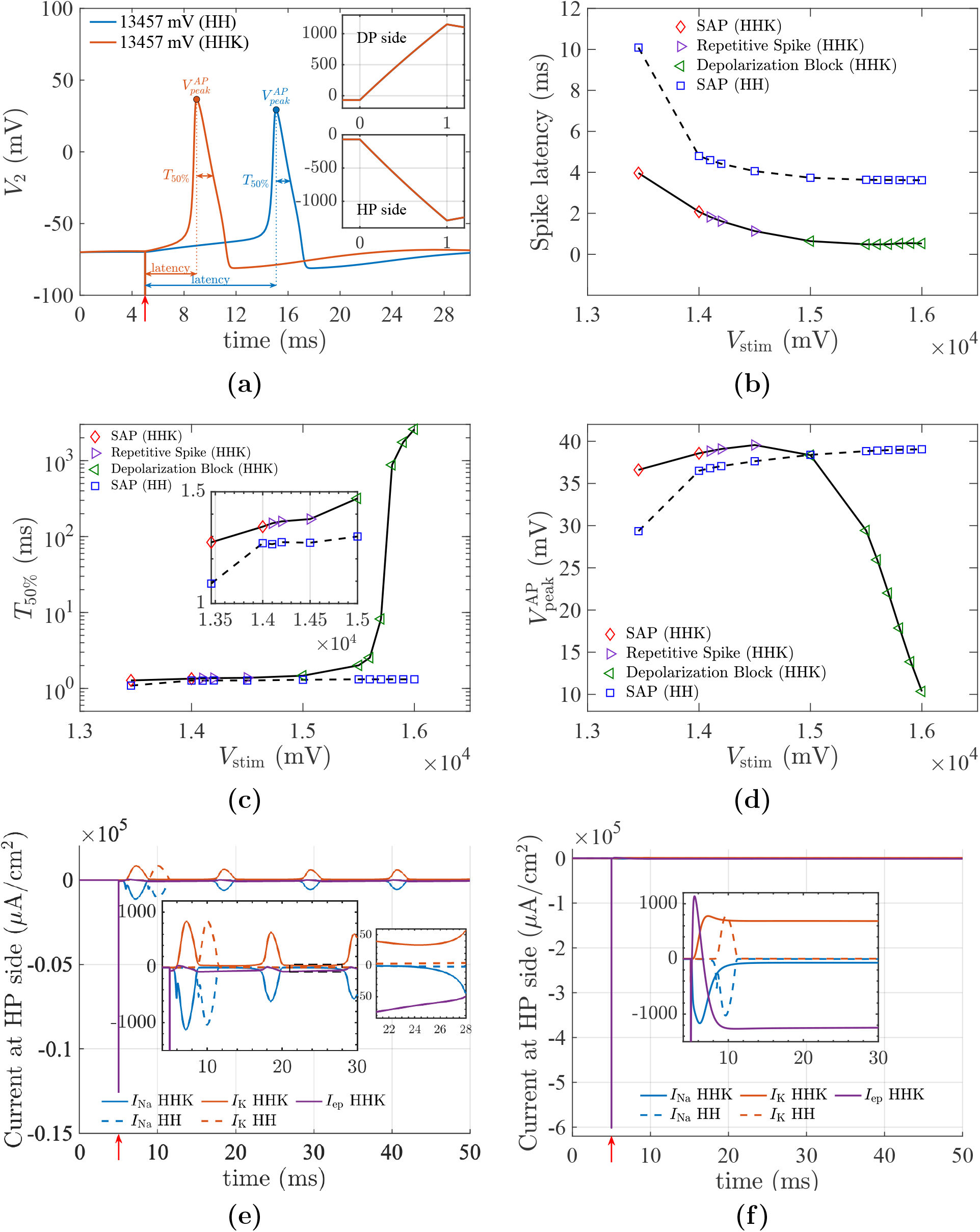
Variation of different model features for a pulse width of *τ* = 1 ns. a) TMP (*V*_2_) for a 1 ns pulse showing single action potentials at *V*_stim_ = 13457 mV. The insets shows capacitive charging at both sides of the membrane (DP and HP side) during the pulse. The time in the inset is reset to *t* = 0 at pulse onset. b) spike latency as a function of *V*_stim_, c) 50% recovery time after spike, & d) Peak TMP during AP vs *V*_stim_ (Solid line - HHK model, dashed line - unporated (HH) model). e) & f) compares ionic and electroporation current in unporated (HH) and HHK models for a 1 ns pulse for *V*_stim_ = 14500 mV and *V*_stim_ = 15500 mV for repetitive spiking and depolarization block respectively. In (e), the smaller inset magnifies the interval between the second and third spikes (21–28 ms, dashed box in the upper inset).The stimulus is applied at *t* = 5 ms, indicated by the upward red arrow(↑).

In depolarisation block, during the pulse, a large number of pores form, which drives a huge electroporation current inside the cell from the HP side as shown in Fig.6(f). However, as *V*_2_ relaxes from a large negative TMP to a slight positive TMP, the electroporation current becomes hyperpolarising during the spiking, hence a decreased amplitude of AP. Sodium current is transient; however, the potassium current stays positive and hyperpolarising, and is balanced by the depolarising electroporation current (see inset Fig.6(f)). The cell reaches back to its RMP once *g*_*ep*_ goes to zero, and that happens when all pores reseal back.

### 3.5 Shorter pulses and smaller neuronal size lower the safety factor

Most electroporation-based treatments are designed to induce pore formation (reversible or irreversible) in biological tissues for cancer therapy. However, this approach is associated with risks of unwanted stimulation. In such applications, it is therefore preferred that electroporation thresholds remain lower than excitation thresholds. In Fig.3(b), a reduced gap between the excitation threshold and the electroporation-driven responses is shown as the pulse duration decreases in the nsPEF regime. For pulses shorter than 20 ns, electroporation of neurons is observed prior to excitation for *RC* = 10^−8^ s, i.e 10 ns (Fig.3(c)). This behaviour is in contrast to that of *RC* = 10^−6^ s, where the relative thresholds remain nearly constant (see inset of the relative SD curve Fig.S19(b)). These results indicate that neurons with smaller RC are more susceptible to electroporative damage in the nsPEF regime.

The variation of thresholds for different cellular responses under a 10-ns pulse across different RC values has been investigated to test this hypothesis. It is known that for short pulses, the TMP 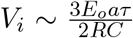] as suggested by the Schwan equation (42). Thus with threshold 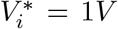, for poration, the *E*_*o*_ required is estimated as 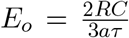] . Since *V*_stim_ can be calculated as 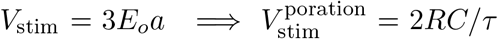] (derivation is provided in the Section S11 of the SI). Thus, for a given *τ*, 10 ns in this case, the required threshold electric field scales as *RC* (Fig.7(a)), while simulations indicated that thresholds scaled exponentially with exponents between 3*/*4 and 4*/*5 (Fig.7(a)), possibly, since it is known that the 1*V* electroporation threshold is not a “hard threshold”. The inset demonstrates that, as *RC* increases, the safety gap between excitation and electroporation is gradually restored (Fig.7(c)).

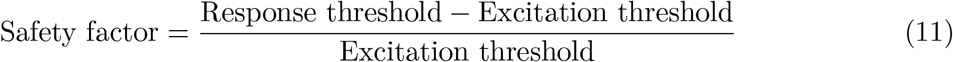

Accordingly, a *safety factor* is defined, which provides a quantitative measure of the relative gap between the excitation threshold and the onset of electroporation-driven responses. The key observations are summarised as follows.

1. The percentage safety factor decreases as the pulse duration decreases for a neuron with *RC* = 10^−8^ s. It falls below -5% for a 1-ns pulse (Fig.7(b)).
2. Pulses in the *µ*sPEF–msPEF regime evoke only a single action potential rather than repetitive spiking or depolarisation block. Although a wide safety gap between excitation and electroporation is restored as the pulse duration increases, the interval between the onset of reversible electroporation (electroporation threshold) and irreversible electroporation (cell death) becomes progressively narrower (see inset of Fig.7(d)).

### 3.6 Mechanism of cell death: NsPEF vs longer pulses

Cell death is defined as the complete loss of a cell’s biological function. It can happen in a programmed way, such as apoptosis, autophagy, or necroptosis, or through external insults that trigger necrosis. In pulsed electric field (PEF) exposure, different pulse durations produce distinct electroporation-mediated routes to cell death. In the present study, nanosecond pulses primarily induce extensive supra-electroporation without irreversible electroporation (IRE), whereas microsecond-to-millisecond pulses predominantly lead to cell death through IRE.

**Fig 7.**
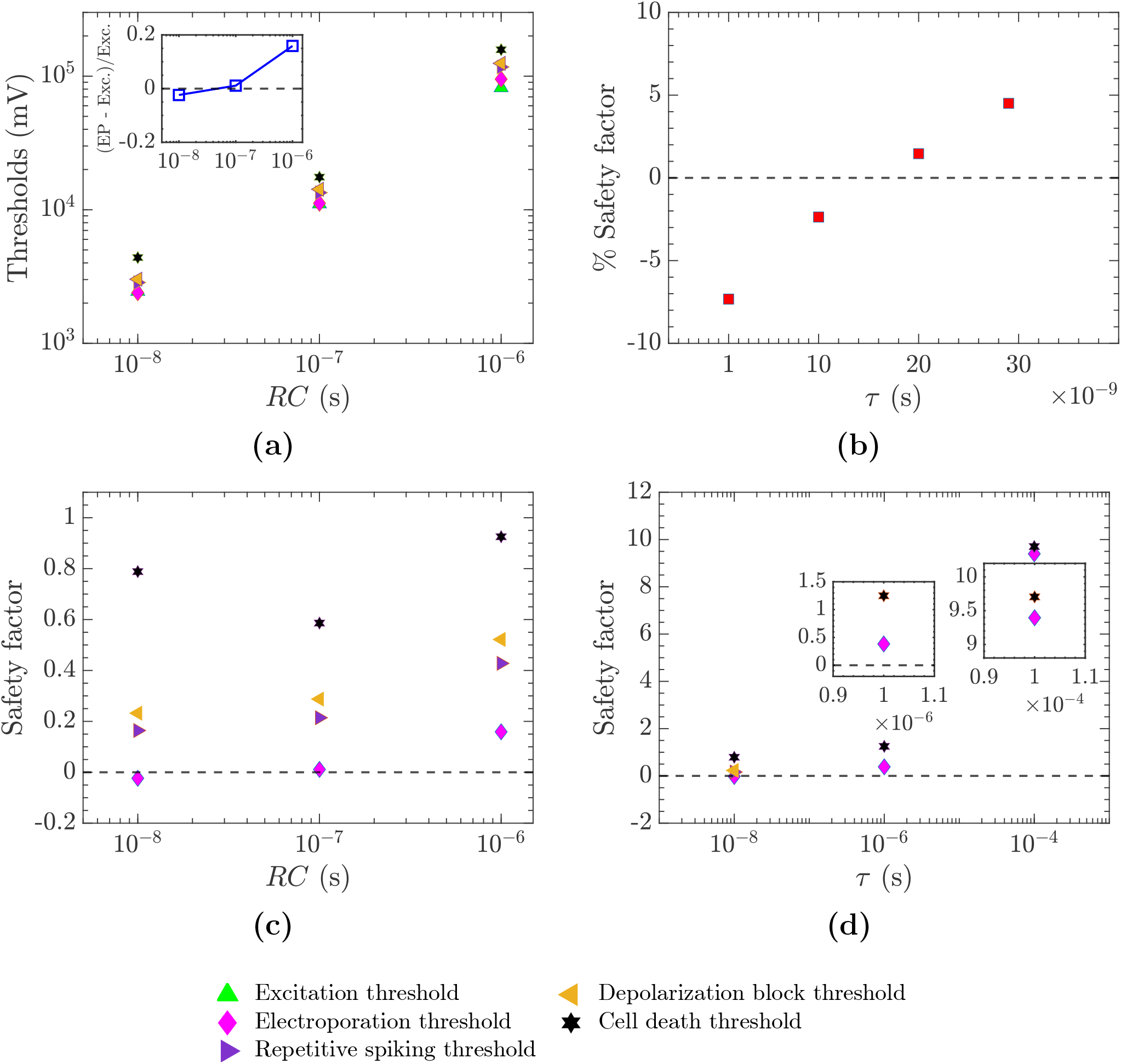
a) Thresholds of cellular responses as a function of *RC* for a 10-ns pulse. The inset shows the relative EP threshold (normalized to the excitation threshold) as a function of *RC*. b) Safety factor (in %) as a function of pulse duration for ultrashort pulses (*<* 30 ns) in a neuron with *RC* = 10^−8^ s. c) Safety factor as a function of *RC* for a 10-ns pulse. d) Safety factor for 10-ns, 1-*µ*s, and 100-*µ*s pulses across different cellular responses in a neuron with *RC* = 10^−8^ s.

Nanosecond pulses admit *supra-electroporation* both in *RC* = 10^−8^ s and *RC* = 10^−6^ s neurons, which is characterized as the formation of a large number of small nanopores. A 10-ns pulse at a *V*_stim_ of 4384 mV is applied to demonstrate this in a *RC* = 10^−8^ s neuron. It causes the formation of a large number of pores both on the DP and HP side with a similar order of magnitude of pores (see Fig.8(a) for HP side, Fig.S20 for DP side). In the pore launching phase, which lasts 20-ns from the onset of stimulus, almost all the large pores are created and launched during the pulse duration marked by the first 10-ns. Out of all the 3595880 large pores formed and launched on the HP side, Pore groups 4 to 10 constitute 3595879 pores, except the pore group 11, which includes a single pore formed just after the pulse, in the relaxation phase. These pores rarely grow beyond nanometer sizes; all of them are absorbed into the small-pore population within the first 150 nanoseconds after the onset of the pulse (see Fig.8(b) including inset, for pore launch during the pulse and absorption into the small-pore population). The sheer abundance of these nanopores leads to a sharp rise in fractional pore area (FPA) during the pore-launching phase (see Fig.8(c)), which in turn causes a sharp fall in membrane tension, as seen in Fig.8(d). These results corroborate the theoretical predictions by Vasilkoski *et al*.(43); a patch of membrane, when subjected to a 10-ns PEF, experiences *supraelectroporation* (a large number of pores whose radii hardly increase), and also reports a similar order of pores created. Previous studies have suggested that electroporation, leading to a fractional pore area, even on the order of ∼ 10^−4^ − 10^−5^ during the pulse, can produce significant physiological effects (44). In the present model, if FPA on either side of the cell exceeds 0.0126 during the pore-launching phase, the membrane tension falls below zero. Under these conditions, IRE is not observed. Instead, the corresponding *V*_stim_ is called as *cell death due to supra-electroporation* (see Appendix C for the limiting FPA calculation). These small pores eventually reseal after an extended period, as seen in Fig.8(e). The small pore density ∼ 10^16^ m^−2^ decays over a time scale ∼ 20 − 30 seconds keeps the TMP depolarised around 0 mV, which eventually reaches the resting potential once all the small pores are resealed, as characterised by the small pore density (*N* ) reaching its equilibrium value, as seen in Fig.8(f). This recovery is prolonged as stimulus strength increases.

In PEFs, another common form of cell death arises due to IRE, which is characterised by the uncontrolled expansion of one or more membrane pores, leading to membrane rupture and loss of cellular integrity. In excitable cells, this process results in a permanent loss of the resting membrane potential, rendering the cell inexcitable and non-functional. IRE is typically associated with the formation of giant, stable pores with radii ranging from approximately 100 nm to 1.5 *µ*m, from which recovery of the resting membrane potential does not occur. We classify this as *cell death due to IRE*, in accordance with several experimental reports (45; 46). Both neurons, *RC* = 10^−8^ s and *RC* = 10^−6^ s, exhibit this form of cell death under *µ*sPEF–msPEF.

Insights into mechanisms of cell death due to IRE are demonstrated here using a 2.9-*µ*s pulse at 1890 mV in an *RC* = 10^−8^ s neuron. There are 10 pores formed during the pulse duration, each in one sub-interval. The first pore launched on the HP side of the cell shows a runaway behaviour with respect to its pore size, as seen in Fig.9(a). The pores launched in subsequent phases decay in size after initial growth and are absorbed into the small-pore population within the first 50 *µ*s after the onset of stimulus. The pore launched in the 1^*st*^ phase continues to grow until the pore size stabilizes at 1.57 *µ*m. It causes the fractional pore area at the HP side to be quite high (∼1.2%)(see Fig.9(b)). Consequently, Membrane tension at the HP side too falls to the order of ∼ 10^−5^ J/m^2^ (0.01 mN/m) and does not recover(Fig.9(c)). It causes the loss of membrane integrity, and the cell does not regain its resting membrane potential of -70 mV (Fig.9(d)). At the HP side, a local minima in FPA (see inset Fig.9(b)) and a local maxima in membrane tension see inset Fig.9(c)) is seen. They feature because all the other pores on the HP side (launched between 2^*nd*^-10^*th*^ phase) are absorbed in the small pore population(*N*_2_) by that time. Similarly, the 3 pores on the DP side are quenched (shown in Fig.S21 in the SI), with minimal increase in fractional pore area (the blip in FPA_1_ and 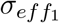 shows the creation, launch and quenching of the three pores at the DP side; see insets Fig.9(b), Fig.9(c)). In general, the DP side sees fewer pores than the HP side, with a difference of several orders of magnitude od pores (compare DP side and HP side pores in Fig.S21). The giant pore causing IRE is usually formed on the HP side.

Experimental evidence also indicates that *µ*sPEF–msPEF pulses can induce cell death via necrosis, pyroptosis, and necroptosis (47). Long-duration pulses favour the growth of smaller pores that expand uninhibitedly. It leads to cell swelling, blebbing, and cell membrane rupture. In the present study, a 2.9-*µ*s pulse at 1890 mV creates a giant stable pore of 1.57 *µ*m on the HP side, resulting in complete loss of resting membrane potential 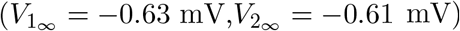]. IRE in the current modelling is consistent with experimental results on necrotic death (48). Furthermore, several experimental studies on nsPEFs have reported apoptotic cell death mediated by mitochondrial pathways and caspase activation (49),(50). Although apoptotic modelling is beyond the scope of the present work, our simulations support the possibility of such a mechanism. NsPEF pulses below 1 *µ*s do not induce IRE, even at higher electric fields.

It creates ∼ 3–4 × 10^7^ pores of 8–9 nm, resulting in a substantial increase in the total porated area. These findings align with experimental reports linking nsPEF exposure to apoptotic outcomes.

## 4 Conclusions

At the outset, the current work aimed to develop a minimal model of an excitable cell that captures the coupled physics associated with excitability and electroporation. Using the model, the interplay between excitation and electroporation is investigated under the regime of nanosecond-pulse electric fields. The model spans the relevant time scales: membrane charging and pore formation on the nanosecond-to-microsecond scale, pore evolution and resealing over microseconds to seconds, and the millisecond-scale responses of the voltage-gated ion channels.

The present computational work suggests that neuronal excitation under nsPEF is governed by the electroporation-mediated shunt conductance. Electroporation was found to enhance conductance predominantly on the hyperpolarised side of the neuron, whereas the depolarised side was affected to a much smaller extent, thereby creating a strong asymmetry in neuronal response. With increasing stimulus strength, excitation response transited from single action potentials to repetitive firing and to depolarisation block due to extensive electroporation. A strength-duration curve highlighting various regimes of excitation and electroporation is also presented. It is shown that an ultrashort pulse (less than or equal to 10 ns) can electroporate neurons without firing an action potential in neurons with small cell-charging time constants (mostly smaller neurons). Although electroporation thresholds, which mark the formation of a single pore on any side of the neuron, do not necessarily result in a significant electrophysiological transition, the altered AP features induced by extensive electroporation may lead to temporary changes in neuronal firing behaviour. In certain cases, it can cause permanent loss of excitability or cell death. While the model does not quantitatively reproduce experimental data, it qualitatively explains key observations reported in the literature. This is because of the obvious differences in the channel dynamics of the expressed voltage-gated ion channels compared with those of the HH model ion channels. The modification of AP features induced by electroporation in the experiments is qualitatively captured by the present model (12). While the current Hodgkin–Huxley–Krassowska model ignores spatial variations in the modelling, it succeeds in elucidating the mechanisms of excitations when electroporation is at play. The current study is significant in shaping new stimulation strategies for neurological and cardiac disorders and in reducing unwanted electrochemotherapy-related pain in cancer patients.

**Fig 8.**
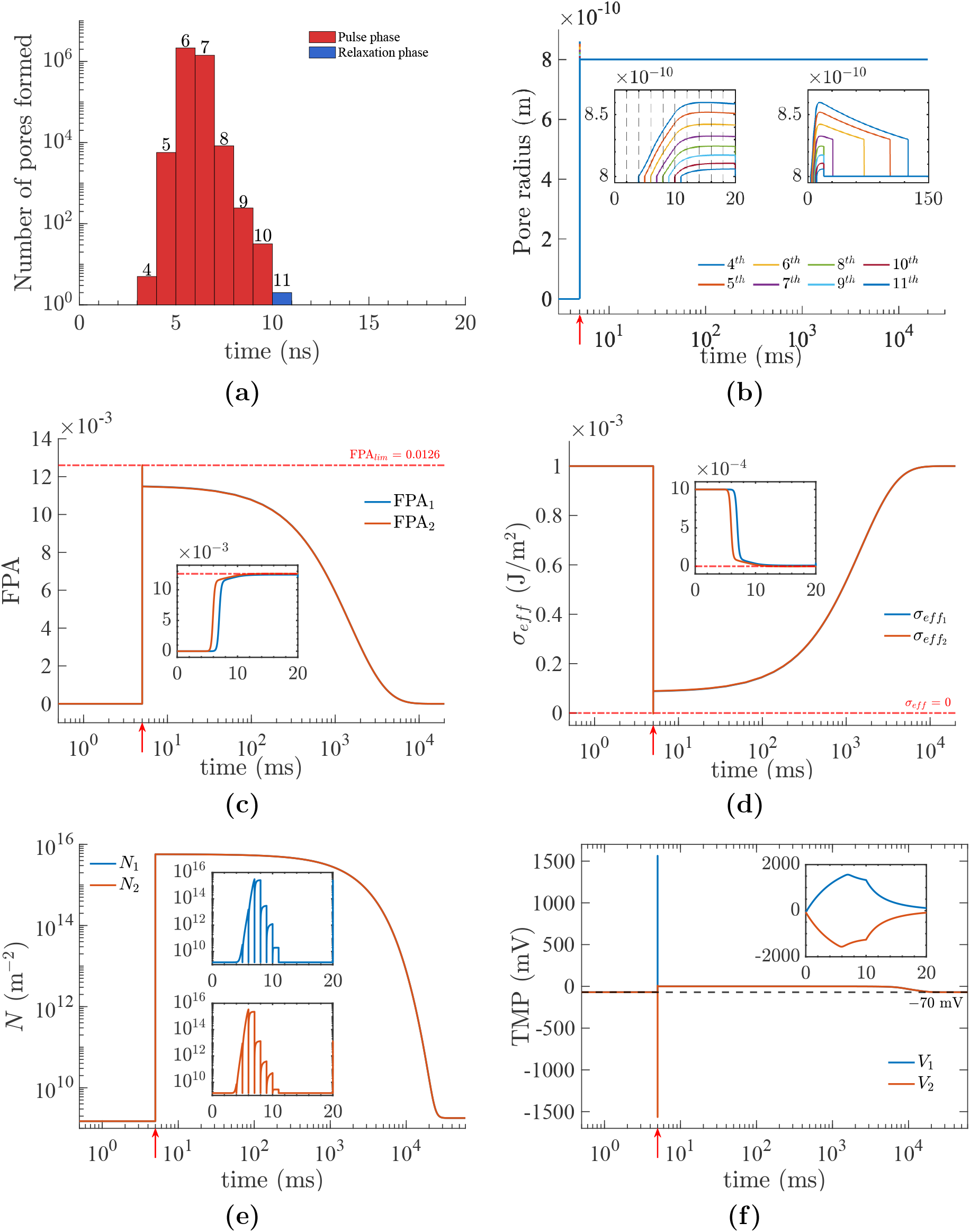
Supra-electroporation induced by a 10-ns pulse with *V*_stim_ = 4384 mV for a neuron with *RC* = 10^−8^ s. (a) Pore statistics on the HP side during the pulse (red bars) and the subsequent relaxation phase (blue bars), with pores initiated over 20 subintervals during the pore formation phase. (b) Corresponding pore radius dynamics for all initiated pores, where the numerical labels denote the corresponding pore initiation subintervals. The left inset enlarges the pore nucleation and growth phase during the pulse, whereas the right inset highlights the longer timescale associated with the resealing of large pores. (c) and (d) Fractional pore area (FPA) and effective membrane tension (*σ*_eff_ ) for the DP and HP sides, indicating the limiting fractional pore area, FPA_lim_, and the corresponding membrane tension. (e) and (f) Small pore density (*N* ) and transmembrane potential (TMP) dynamics, respectively. The insets show the pulse-duration dynamics with the time axis reset to *t* = 0 at pulse onset and displayed on the nanosecond timescale for improved visualization. The stimulus onset at *t* = 5 ms is indicated by the upward red arrow(↑).

**Fig 9.**
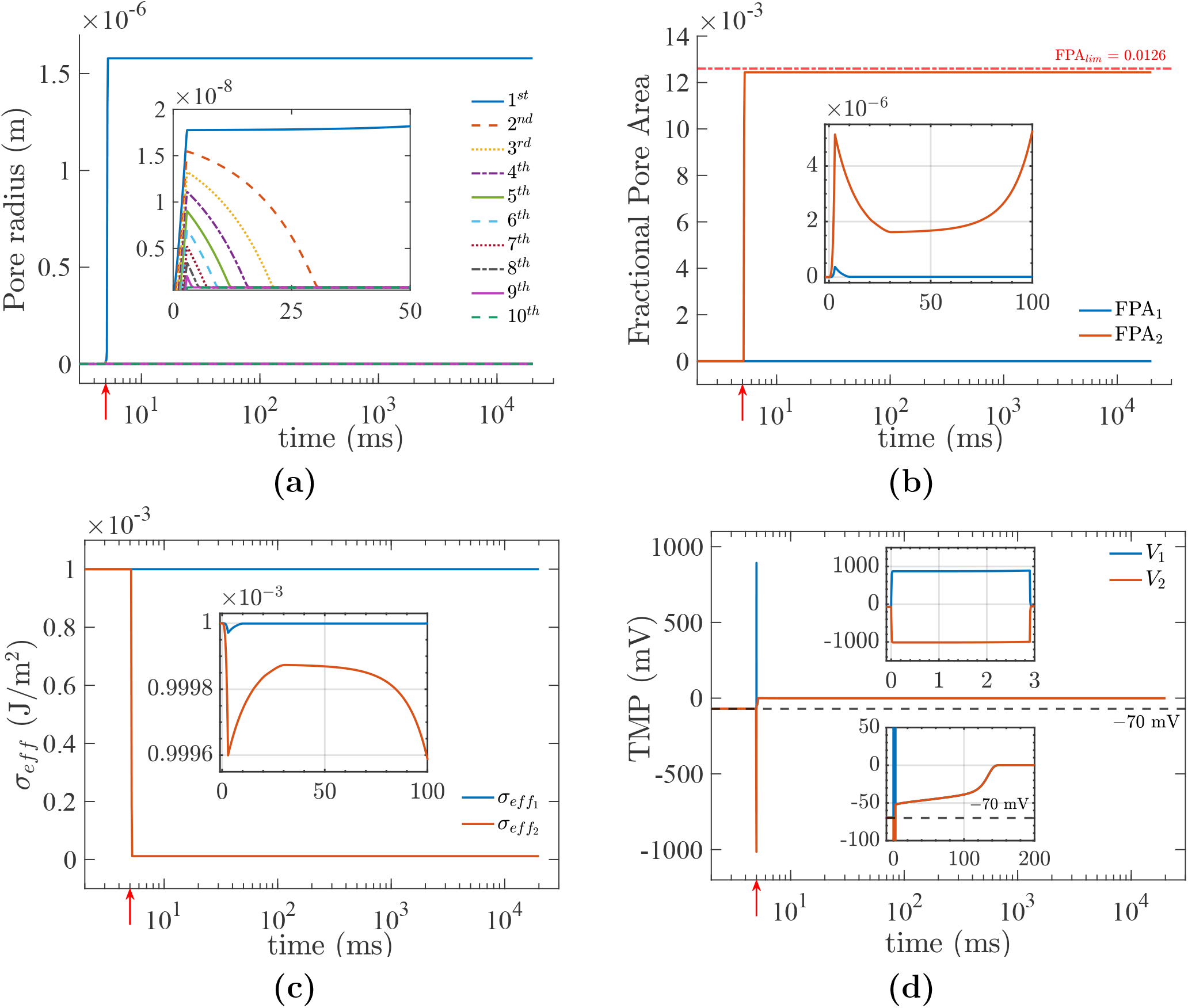
Irreversible electroporation (IRE) induced by a 2.9 *µ*s pulse with *V*_stim_ = 1890 mV for a neuron with *RC* = 10^−8^ s. (a) Pore radius dynamics on the (HP) side for 10 pores initiated at different time instances, labeled 1–10 to indicate their order of formation. The inset highlights the time window during which pores nucleate, grow, and reseal, except for the dominant (giant) pore. (b) Fractional pore area and (c) membrane tension dynamics on the DP and HP sides, respectively. (d) Transmembrane potential (TMP) dynamics; the time axis in the insets is reset to *t* = 0 at pulse onset and displayed on the microsecond (*µ*s) timescale for improved visualization. The stimulus onset at *t* = 5 ms is indicated by the upward red arrow(↑).

**Fig 10.**
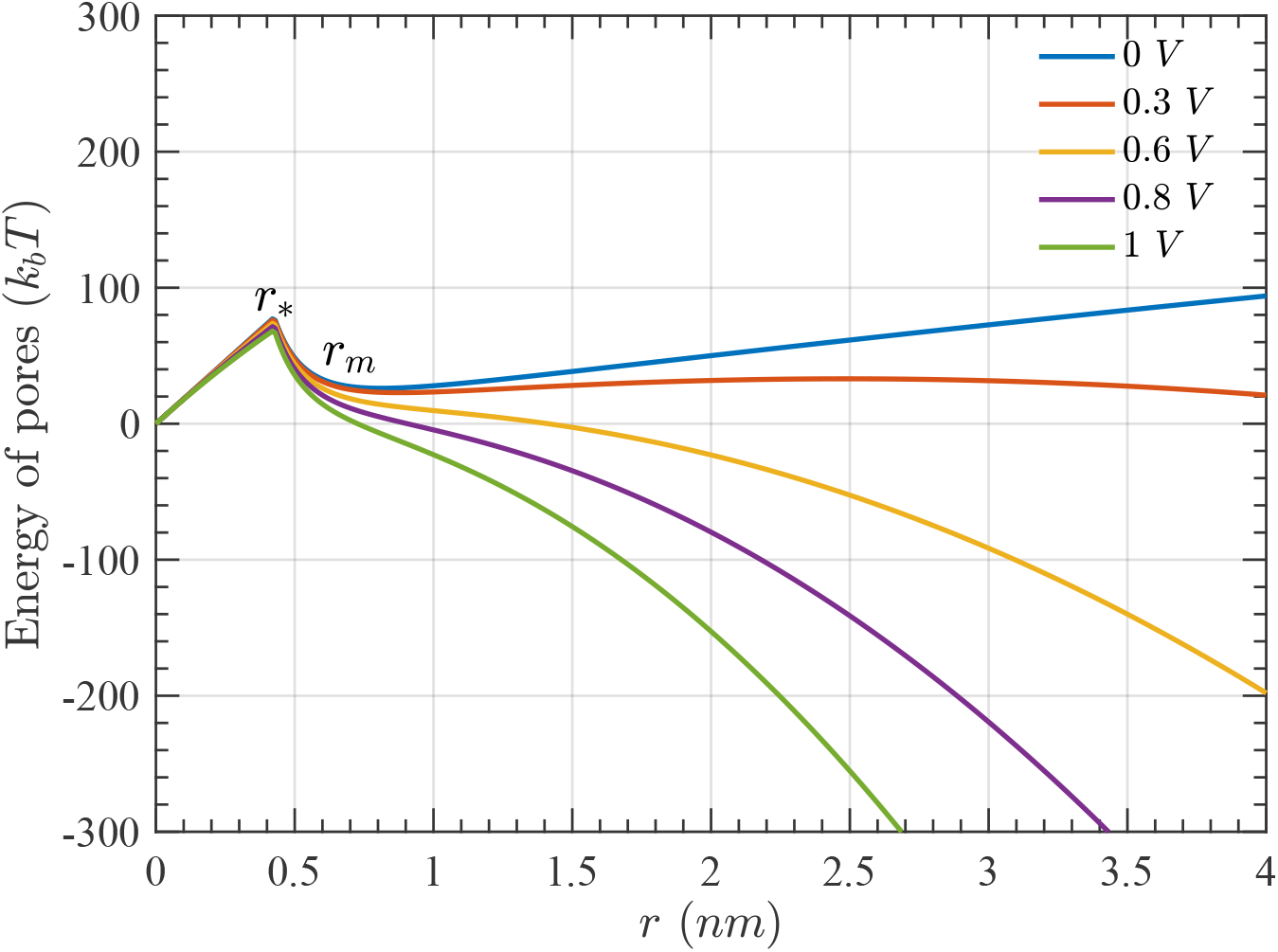
Free energy of a pore as a function of pore radius for different values of the transmembrane potential (TMP).

## Appendix

### A Electroporation Model (ASE)

The asymptotic analysis of the continuum model of electroporation, the Smoluchowski Equation (SE), is a landmark work in the domain of electroporation modelling. It reduces the computationally intensive PDE to a set of ODEs, which are more tractable. It is known as the Asymptotic Smoluchowski Equation (ASE), details of which can be found in (17; 22), and are also briefly described here:

1. Energy of pores as a function of their radius describes the existence of two kinds of pores - hydrophobic (non-conducting) and hydrophilic (conducting) pores. For a given membrane potential *V*_*m*_, there exist energy barriers corresponding to radii *r*_∗_ and *r*_*d*_. Hydrophobic pores *r < r*_∗_, if formed, are quickly destroyed by lipid fluctuations. However, pores of radii *r* ≥ *r*_∗_ are hydrophilic pores. These pores expand to an energy minima to the right of *r*_∗_. In the context of the present work, all pores are assumed to be created hydrophilic with radius *r*_*m*_.
2. Kinetics of the formation of these pores is described by the equation,

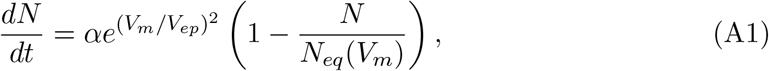

where *N* is the pore density, *V*_*m*_ is the TMP, *V*_*ep*_ is the characteristic voltage of electroporation, and *α* is the creation rate coefficients. *N*_*eq*_ is the equilibrium pore density at a given TMP, *V*_*m*_.

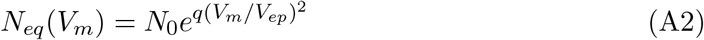

*N*_0_ is the equilibrium pore density at zero TMP and *q* = (*r*_*m*_*/r*_∗_)^2^.

1. These hydrophilic pores grow in the pore-radius space. The governing equation for the evolution of K pores formed is given by,

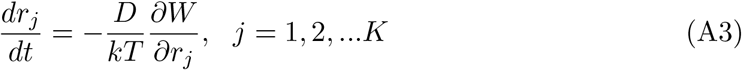

where *D* is the diffusion coefficient of the pore radius, *k* is the Boltzmann constant, and *T* is the absolute temperature. *W* is the bilayer energy,

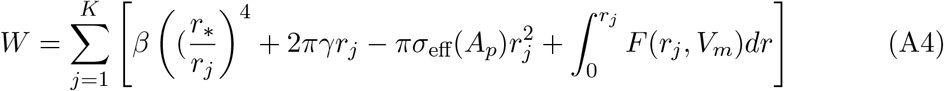

The first term represents steric repulsion between lipid heads, the second term is the edge energy cost because of the pore perimeter, the third term represents the pore energy released due to membrane tension, and the fourth term is the electrical energy due to TMP. *σ*_eff_ is calculated using the area of the pores formed. The pore area is calculated as 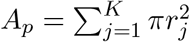].

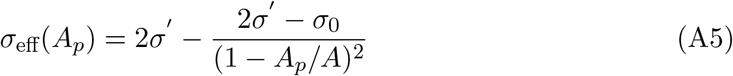

*σ*_eff_ is the effective tension, whereas *σ*_0_ is the membrane tension without pores. A is the total area of the lipid bilayer membrane. *σ* ^*′*^ is the energy per unit area of the hydrocarbon-water interface. *σ*_eff_ decreases as *A*_*p*_ increases. It couples the membrane tension across the pores. As a result, each pore experiences a decrease in membrane tension. The following equation gives the electrical energy as a result of electroporation,

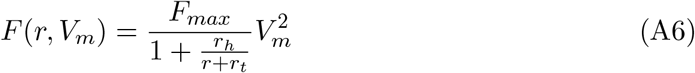

*F*_*max*_, *r*_*h*_, and *r*_*t*_ are constants (detailed in section S1.1 of the SI). The transmembrane potential *V*_*m*_ is calculated and updated using an electrical model. In the current study, a two-layer membrane patch coupled via cytoplasmic resistance is considered, and a corresponding equivalent electrical circuit is provided in the methodology.

### B Numerical Implementation Details

The model described by Eqs. 1 to 8, as well as Eqs. A1 - A6 is implemented using MATLAB. The numerical integration of Eqs. 1, 2, 4 and A3 is done using an in-built solver *ode23s*, which is suited for stiff systems and provides computational efficiency for the voltage-dependent gating and electroporation dynamics. The total integration time (*t*_*f*_ ) is set to 20 s. In the numerical simulations, a monopolar rectangular pulse is given at *t* = 5 ms. This ultrashort pulse of high intensity, typically, is on the order of -ns to -*µ*s. The pulsing can result in the formation and evolution of conducting (hydrophilic) pores. This happens over a time scale of a few -*µ*s to ns. However, excitation or firing of AP in neurons happens over few *milliseconds* . Resealing of these hydrophilic pores happens through their conversion into hydrophobic (non-conducting) pores and eventual annihilation by the lipid fluctuations. This occurs on a time scale of a few *seconds*.

To efficiently capture these multi-scale, multiple phenomena, the integration time is divided into four sub-time spans, namely *t*_1_, *t*_2_, *t*_3_, and *t*_4_. The *equilibrating phase* (*t*_1_), which begins at *t* = 0 and lasts until *t* = 5 ms, ensures that all the state parameters are at steady state before the pulse. *t*_2_ is the duration of the pulse, while *t*_3_ equals *RC* of the cell, the timescale over which the TMP relaxes due to cytoplasm resistivity. The magnitude of the membrane potential at both ends increases during the pulse, and the recovery post-pulse to the resting potential occurs over a few *time constants* of the cell. The combined period of *t*_2_+*t*_3_ is the phase in which pores form and evolve. This phase is called *pore launching phase*. At the end of this phase, the cell reaches near its resting potential, which is well below the electroporation threshold. The final integration phase *t*_4_ absorbs all the shrunken pores. Hence, it can be termed as *pore absorbing phase* (Pore nucleation, evolution and absorption strategies are detailed in Section S13 of the SI). Each of these subdivisions requires a distinct solver configuration.

The model assumes two pore sub-populations to represent two kinds of conducting pores at both DP and HP ends of the membrane. All pores are created as small pores with radius *r*_*m*_ = 0.8 nm. Their population is represented by pore density *N* (*t*), whose dynamics is described by Eq. 4. These small pores, once formed, are allowed to grow and are called large pores. The large pores (*r > r*_*m*_) are considered individually and evolve as per Eq. A3. Initially, the membranes at both ends are at resting membrane potentials of −70 *mV* . At *t* = 0, gating variables at both the ends are *m*_0_ = 0.053, *n*_0_ = 0.317 and *h*_0_ = 0.596. The small pore densities, *N*_1_ and *N*_2_, equal *N*_0_ and it is assumed that there are no large pores. The flow of a typical simulation time loop is as follows:

1. *Equilibration phase*: During the *equilibrating phase*, no external stimulus is applied. In this phase, only 10 ode’s get integrated, namely the membrane potential (*V*_*i*_), their corresponding ion-channel gating variables (*m*_*i*_, *n*_*i*_, *h*_*i*_) and small pore densities (*N*_*i*_), *i* = 1, 2. This just equilibrates the system to a consistent steady state before the stimulus. As no rapid dynamics are expected, default time steps and tolerances are used, allowing for longer adaptive time steps and efficient convergence.
2. *Pore launch phase*: The applied pulse is turned on for the duration *t*_2_. This causes the membrane potential to rise up exponentially fast. After the pulse is switched off, the membrane potential relaxes over *t*_3_. Both of these phases involve electroporation since the TMP is much above 1 V. Hence, pore creation and launching modules are invoked. These are based on Smith *et al*.[2004] with some modifications and are also explained below for the sake of completeness(22).
  a. Both *t*_2_ and *t*_3_ are further subdivided for computational accuracy. It was found that ten subdivisions in each *t*_2_ and *t*_3_ yield reasonable results, although finer subdivisions can be considered. For the first subdivision, *V*_1_, *V*_2_, *m*_1_, *m*_2_, *n*_1_, *n*_2_, *h*_1_, *h*_2_, *N*_1_ and *N*_2_ are solved for.
  b. Eqn.4 is used to evaluate *N*_1_ and *N*_2_ after each subdivision. An increase in *N* means the creation of small conducting (hydrophilic) pores, while a decrease in *N* means their annihilation into non-conducting (hydrophobic) pores.
  c. The minima (*r*_*m*_)) in the energy landscape Fig.10 disappear due to the applied voltage to the membrane. This allows the small pores to grow. After every subdivision, *floor*(*N* (*t*)*A*) is the consistent number of large pores that are launched by equivalent concession of small pores and transferred into the large pore population described by *K*. However, if *floor*(*N* (*t*)*A*) = 0, it indicates that no large pores are launched in that subdivision. The values of *N* is updated accordingly. To reduce computational cost, instead of tracking the growth of individual pores, all pores formed in a given subdivision are treated as a single group of large pores. These pores are initialized with a radius of *r*_*m*_ = 0.8 nm at the time of launching and evolve collectively. This procedure is applied to both the depolarized (DP) and hyperpolarized (HP) sides of the membrane.
  d. Each subdivision sees the evolution of the radii of large pore groups along with the membrane potentials (*V*_1_ and *V*_2_), ion channel-gating variables (*m, n,h*), and the small pore densities (*N*_1_ and *N*_2_). Solving for membrane potential requires calculation of shunt conductances (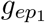] and 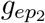] ) which are calculated using the dynamic pore area (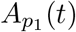] and 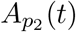]). The membrane tension (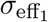] and 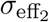] ) are also evaluated dynamically during each subdivision. All these constitute a self-consistent system of ODEs, which is solved using the *ode23s* solver.
  e. During the *t*_2_-phase, ultrashort pulses lead to the deactivation of potassium channels and the de-inactivation of sodium channels. The large negative TMP attained at the hyperpolarized side during nsPEF stimulation causes immediate deactivation of K-channels and de-inactivation of Na-channels on that side. The resulting decrease in potassium efflux, combined with an increase in sodium influx, continues to depolarize the membrane. As the system transitions into the *t*_3_-phase, the TMP begins to relax, and the gating variables *n*_2_ and *h*_2_ continue to evolve based on both the TMP and the injected current, during and after the stimulus. Capturing these rapid transients necessitates careful selection of the time step (Δ*t*) during integration in the pore launching phase.
  f. Due to the rapid transients in TMP, ion-channel dynamics and electroporation dynamics during this phase, smaller time steps are essential for numerical stability and accuracy. A strict upper bound on the time step (Δ*t*) is imposed via odeset to accurately capture fast electroporation kinetics and ion-channel dynamics. In the *t*_2_-phase, for nanosecond pulsed electric fields (nsPEFs), the maximum step size is set to 10^−11^ s, whereas for microsecond pulses (*µ*sPEFs), it is relaxed to 10^−8^ s. In the *t*_3_-phase, where membrane voltage relaxes over the *RC*-timescale, the maximum step size is made *RC*-dependent. For example, for cells with *RC* = 10^−5^ s a maximum time step of 10^−11^ s is used, while those with *RC* = 10^−6^ s are assigned a maximum step size of 10^−9^ s.
  g. To account for the possibility of irreversible electroporation during the pore launching phase, the simulation actively monitors the pore radii. A threshold of 100 nm is defined for irreversible electroporation; if the radius of any pore group exceeds this threshold, the simulation is terminated, and the cell is deemed to have “irreversibly electroporated”.
3. *Pore absorption phase*: The end of the pore launching phase marks the recovery of the membrane potential toward its resting value. In the *pore absorption phase*, large pore populations are absorbed into the small pore population if the radius of any pore group falls below the absorption threshold *r*_abs.th_, which is set to 0.83 nm. This is implemented by removing the corresponding pore group from the large pore population *K*, and updating *N* accordingly. This absorption process occurs on a fast timescale, typically within microseconds. Once all large pore groups are absorbed, the small pore densities *N*_1_ and *N*_2_ relax over a much slower timescale of several seconds. This phase, denoted as *t*_4_, continues until the end of the simulation at time *t*_*f*_ . It is during this phase that the action potential is typically observed.

As the small pore densities *N*_1_ and *N*_2_ evolve slowly while the system returns to baseline, the time-step restrictions are relaxed to their default values, since this phase does not exhibit numerical stiffness. The ion channel dynamics and pore recovery processes in this phase occur over milliseconds, allowing the solver to adapt more freely while maintaining stability in the slow resealing dynamics. However, the error tolerance limits (both absolute and relative) should be carefully chosen based on the desired accuracy. Conservative tolerance settings are recommended to ensure numerical reliability of the results.

The code is rigorously benchmarked in the two limits of an unporated neuron (Boinagrov *et al*.) and a porated, unexcitable cell (Smith *et al*.) (Sections S1, S3 and S4, see Figures S2, S5, S6 and S7).

**Table A1.** Time constants of the Hodgkin–Huxley gating variables (*m, n*, and *h*) at physiological transmembrane potentials (TMPs) and at peak TMPs attained on the depolarizing (DP) and hyperpolarizing (HP) membrane sides during a 100-ns pulse for a neuron with *RC* = 10^−8^ s. The peak TMPs correspond to applied stimulus amplitudes *V*_stim_ = 1557, 2100, 2300, and 2479 mV. All time constants are expressed in milliseconds.

| TMP (mV) | $\tau_m$ | $\tau_n$ | $\tau_h$ |
| --- | --- | --- | --- |
| **Physiological Range** |  |  |  |
| −70 | 0.2368 | 5.4586 | 8.5160 |
| −50 | 0.4790 | 3.9132 | 3.3934 |
| 0 | 0.2159 | 1.5300 | 1.0161 |
| +40 | 0.1175 | 0.9693 | 1.0000 |
| **Peak TMPs (DP Side)** |  |  |  |
| +711 | $1.32 \times 10^{-2}$ | 0.1297 | 1.0000 |
| +980 | $9.80 \times 10^{-3}$ | $9.62 \times 10^{-2}$ | 1.0000 |
| +1081 | $8.90 \times 10^{-3}$ | $8.76 \times 10^{-2}$ | 1.0000 |
| +1180 | $8.20 \times 10^{-3}$ | $8.06 \times 10^{-2}$ | 1.0000 |
| **Peak TMPs (HP Side)** |  |  |  |
| −850 | $3.78 \times 10^{-20}$ | $4.66 \times 10^{-4}$ | $1.65 \times 10^{-16}$ |
| −1193 | $2.00 \times 10^{-28}$ | $6.40 \times 10^{-6}$ | $5.88 \times 10^{-24}$ |
| −1219 | $4.73 \times 10^{-29}$ | $4.63 \times 10^{-6}$ | $1.60 \times 10^{-24}$ |
| −1308 | $3.38 \times 10^{-31}$ | $1.52 \times 10^{-6}$ | $1.87 \times 10^{-26}$ |

### C Cell death due to supra-electroporation

Pore formation is always accompanied with the reduction of bilayer energy, which equals membrane tension times pore area However, if multiple pores form and expand simultaneously, the membrane tension doesn’t remain constant It decreases with the increase of total pore area The Equation A5, calculates the effective membrane tension during pore formation, evolution, and resealing However, in the case of extremely high nsPEF voltages, the cell membrane experiences huge electroporation or supraelectroporation creating ∼ 10^7^ pores of 0.8 nm These huge numbers of pores formed in the pore launching phase, push the membrane tension beyond its limits. The effective membrane tension values must be positive to ensure the stable existence of pores. This put a limit on the fractional pore area, FPA, which is defined as the ratio of pore area by the membrane area for a given side of the membrane It is calculated at both side of the membrane The membrane tension must be positive, as described below:

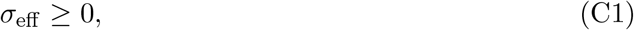

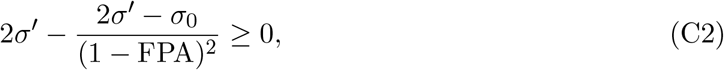

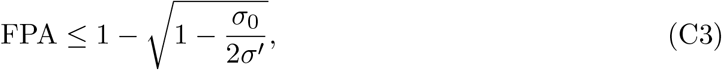

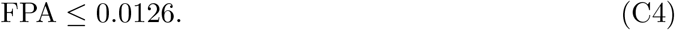

For FPA *>* 0.0126, the stability of pores pushes the area of the cell *A*. This can cause necrotic cell death.

## Data availability

All data and code needed to reproduce the findings of this study are publicly available. The MATLAB implementation of the Hodgkin–Huxley–Krassowska (HHK) model, the single-membrane electroporation model used for benchmarking, the saved simulation outputs, and the scripts that generate every figure and table in the manuscript and Supporting Information are available at https://github.com/rashidcompneuro/HH-Electroporation-PLOSCB and are archived at Zenodo (DOI: 10.5281/zenodo.23409120). The repository is released under the MIT License. Benchmark data digitised from Smith *et al*. (2004) and Boinagrov *et al*. (2010) are included in the repository. All other relevant data are within the manuscript and its Supporting Information files.

## Acknowledgments

The results are generated using the numerical code in MATLAB. This code is an in-house development without any assistance from LLMs. The use of generative AI (Claude, Anthropic) was restricted to documenting the public code repository and improving the grammar and clarity of selected portions of the manuscript, which were reviewed and verified by the authors, who take full responsibility.

## Supporting information

**S1 Text. Supplementary information**. Benchmarks of the electroporation and HH models, numerical convergence, pore and ionic-current dynamics, scaling laws and derivations (Sections S1–S16, Figs S1–S22, Tables S1–S7).

